# Magnetically guided macrophage immunobots coordinate iron-metabolic cascades and immunogenic ferroptosis for tumor immunotherapy

**DOI:** 10.64898/2026.05.29.728625

**Authors:** Xirui Zeng, Yingfei Zhang, Chong Wu, Qijia Duan, Mengqi Fan, Kuan Wang, Shirong Cai, Immihan Ceren Yasa

## Abstract

Adoptive cell therapy (ACT) remains challenging in solid tumors, where poor tumor infiltration, metabolic heterogeneity, and an immunosuppressive tumor microenvironment (TME) constrain therapeutic efficacy. Here, we developed a magnetically actuated macrophage-based immune microrobot (immunobot) for active solid tumor immunotherapy. Immunobots were constructed by loading bone marrow-derived macrophages (BMDMs) with lipopolysaccharide-modified Janus L1₀-FePt magnetic microrollers (LMRs), enabling hard-magnetic actuation. Optimized LMR loading supported robust propulsion, retention under flow, and enhanced barrier penetration. LMRs further promoted M1-like polarization through LPS-driven inflammatory activation and FePt-derived labile iron-amplified oxidative stress, with altered iron homeostasis, increased reactive oxygen species (ROS), and enhanced NF-κB signaling. Immunobots also induced ferroptosis-associated immunogenic cell death in tumor cells. *In vivo*, magnetically guided immunobots suppressed tumor growth, reprogrammed tumor-associated macrophages (TAMs), promoted dendritic cell maturation, and enhanced CD8⁺ T cell activation. This work establishes a microrobotic immunotherapy platform for active magnetic transport, iron-metabolic regulation, and immune remodeling in solid tumors.

## Introduction

Adoptive cell therapy (ACT) harnesses living immune cells to recognize and eliminate malignant cells and has transformed the field of cancer immunotherapy (*1–3*). Although T cell–based ACT has achieved remarkable clinical success in hematologic malignancies, its broader application to solid tumors remains limited by multiple barriers within the tumor microenvironment (TME), including antigen heterogeneity, limited cellular persistence, functional exhaustion, immunosuppressive signaling, and inadequate tumor infiltration (*3*). Thus, alternative immune effector cell platforms are needed to overcome these limitations and improve therapeutic outcomes in patients with advanced solid tumors.

Macrophages are innate immune cells with marked phenotypic plasticity that are abundant in many solid tumors and respond to inflammatory mediators, chemokines, and hypoxic cues within the TME, supporting their natural recruitment to tumor tissues, including poorly vascularized hypoxic regions (*4–6*). Beyond their intrinsic tumor-homing capacity, macrophages mediate phagocytosis, antigen presentation, cytokine secretion, and extracellular matrix remodeling, thereby coordinating innate and adaptive immune responses (*6–8*). However, tumor-associated macrophages (TAMs) frequently acquire tumor-supportive and immunosuppressive states that promote tumor growth, angiogenesis, metastasis, and therapeutic resistance, contributing to poor patient outcomes (*7*, *8*). To harness macrophages therapeutically, engineered macrophage strategies, including chimeric antigen receptor macrophages (CAR-Ms) and macrophages equipped with cytokine-releasing cellular backpacks, have been developed to enhance tumor recognition, phagocytic clearance, and sustained proinflammatory activity within solid tumors (*9*, *10*). Nevertheless, following systemic administration, the therapeutic efficacy of these strategies remains dependent on effective tumor trafficking, retention, and tissue distribution. Therapeutic macrophages must withstand hemodynamic shear stress, undergo vascular arrest and endothelial adhesion, and complete transendothelial migration. After extravasation, dense extracellular matrices, elevated interstitial pressure, and aberrant tumor mechanics further restrict deep tissue infiltration and spatially uniform distribution (*2*, *11*, *12*). These limitations highlight the need for biohybrid platforms that preserve macrophage-mediated immune functions while introducing externally controllable capabilities for active navigation, vascular retention, and enhanced tumor infiltration.

Externally powered propulsion strategies for biomedical microrobots include optical, acoustic, electrical, and magnetic actuation (*13*). Among these, magnetic guidance is particularly attractive for living cell–based microrobots because magnetic fields penetrate deep tissues with minimal attenuation, are compatible with biological environments, and enable remote spatiotemporal control over motion and retention (*13*, *14*). For controlled transport under flow, Janus microrollers provide an effective architecture in which structural asymmetry converts externally applied magnetic torque into directional rolling along biological boundaries (*15*). Their propulsion performance critically depends on the magnetic material, which governs torque generation and dynamic stability under rotating magnetic fields. Chemically ordered, face-centered tetragonal L1₀-FePt is an attractive hard magnetic material because of its high magnetocrystalline anisotropy and remanence, enabling strong magnetic actuation, biocompatibility and improved performance under flow relative to previously used soft magnetic or nickel-based microroller systems (*16*). Consistent with this design principle, macrophage-based microrobots incorporating FePt/LPS-functionalized Janus particles have been explored for remote magnetic actuation, imaging, and antitumor activity (*17*). Thus, equipping macrophages with hard magnetic FePt microrollers provides a promising strategy for combining active mechanical navigation with intrinsic tumor-homing and immune functions.

Beyond enabling magnetic actuation, iron-containing magnetic materials may provide additional therapeutic functions in biohybrid systems. For example, superparamagnetic iron oxide nanoparticles can be internalized by macrophages and, under specific conditions, promote proinflammatory and antitumor phenotypes, consistent with polarization-associated differences in macrophage iron handling (*18–20*). Separately, FePt-based nanomaterials have been shown to promote reactive oxygen species accumulation and lipid peroxidation in tumor cells, thereby inducing ferroptosis (*21*, *22*). Ferroptosis is a regulated cell death modality driven by iron-dependent lipid peroxidation (*23*). Under specific conditions, ferroptotic tumor cells can expose pro-phagocytic and immunostimulatory signals, promote macrophage-mediated clearance, and engage adaptive antitumor immune responses, although these immunological consequences are context- and stage-dependent (*24–27*). Despite these advances, whether a macrophage-based FePt microrobotic platform can integrate active transport with ferroptosis-associated immunogenic tumor injury and subsequent immune remodeling *in vivo* remains to be elucidated.

In this study, we developed a macrophage-based magnetically controlled biohybrid immunorobot (Fig.1). The system comprises bone marrow-derived macrophages (BMDMs) loaded with lipopolysaccharide (LPS)-modified cationic L1₀-FePt microrollers (LMRs), thereby integrating magnetic field driven active navigation with macrophage immune reprogramming for solid tumor immunotherapy. We utilized LMRs as both a magnetically responsive, actuator unit and a biochemical and ferroptosis-inducing regulator. With optimized intracellular loading, LMRs enhanced the magnetically controlled motility and transendothelial migration of macrophages, while LPS-mediated proinflammatory stimulation together with FePt-associated iron-dependent oxidative stress promoted and sustained M1-like polarization. The immunobots induced ferroptosis-associated immunogenic injury in tumor cells, characterized by glutathione (GSH) depletion, glutathione peroxidase 4 (GPX4) downregulation, and lethal lipid peroxidation (LPO) accumulation. *In vivo*, magnetically guided immunobot treatment suppressed tumor growth and was accompanied by an increased intratumoral M1/M2 macrophage ratio, enhanced dendritic cell maturation, and activated CD8⁺ T cell responses. Together, this platform integrates active delivery, macrophage immune reprogramming, ferroptosis-associated immunogenic tumor injury, and adaptive immune activation within a single biohybrid system, offering a promising strategy for cell-based immunotherapy of solid tumors.

**Fig. 1.**
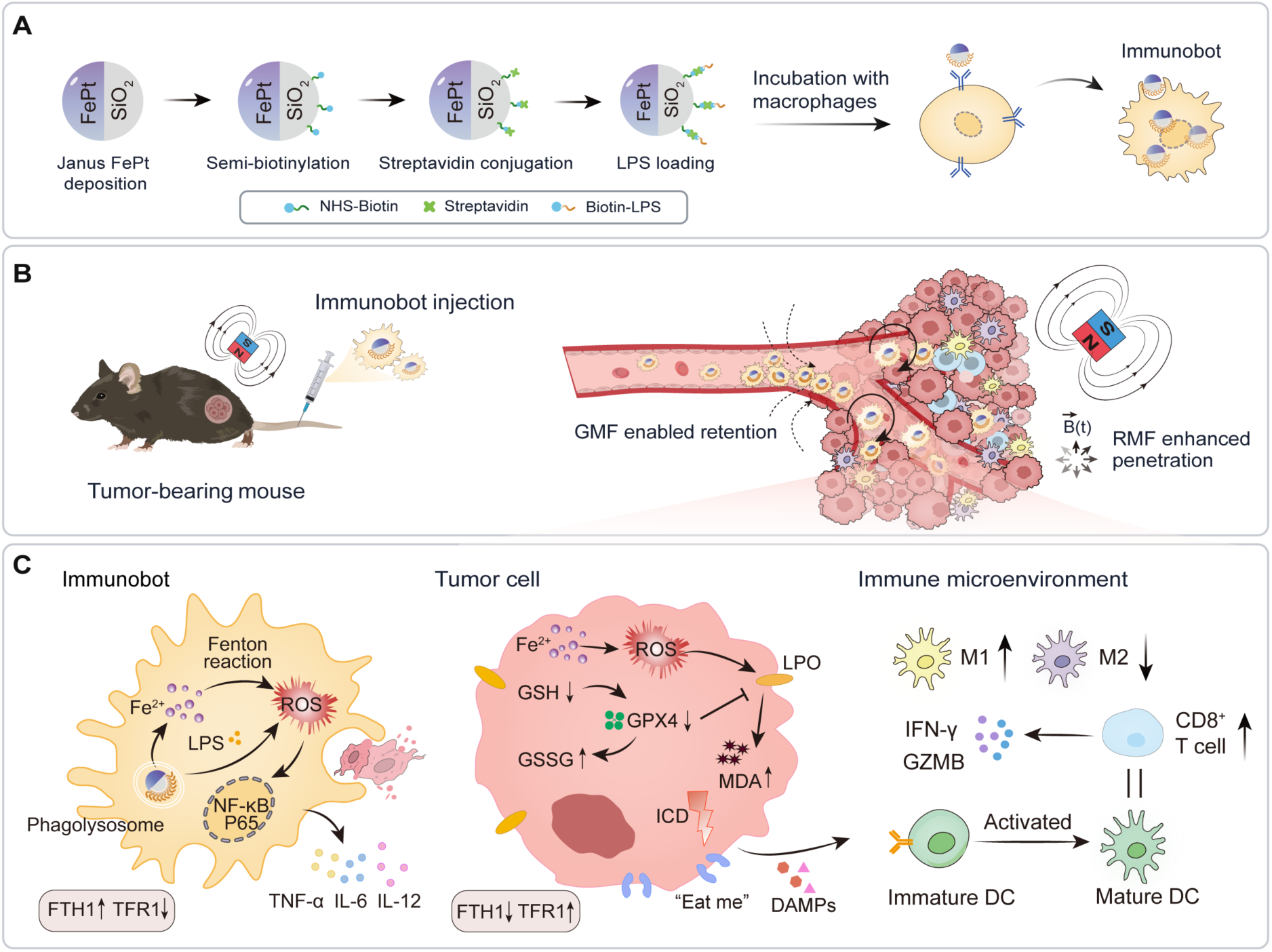
Construction and therapeutic mechanism of immunobots. (**A**) Preparation of LPS-loaded magnetic immunobots through sequential surface functionalization of Janus FePt/SiO₂ particles and incubation with macrophages. (**B**) Immunobots realizing enhanced tumor accumulation and penetration under hybrid magnetic field application. (**C**) Internalized LMRs promote M1-like polarization of immunobots through LPS-mediated inflammatory activation and FePt-derived iron-amplified oxidative stress, while inducing ferroptosis-associated ICD in tumor cells to activate antitumor immunity and remodel the immunosuppressive TME.

## Results

### Immunobot magnetic and biochemical unit design with payload-dependent heterogeneity

To generate robust magnetically actuated Janus microrollers, 5 µm SiO₂ microspheres were hemispherically coated with FePt by direct current (DC) magnetron co-sputtering using independent Fe and Pt targets (Fig. 2A). In contrast to monoatomic-layer molecular beam epitaxy deposition that has commonly been used for L1_0_-FePt films at sub-Å/s deposition rates, we optimized this route as a rapid thin-film deposition strategy for batch-scale fabrication. Considering the hard-magnetic response of FePt arises from chemically ordered, near-equiatomic L1_0_ phases, and magnetocrystalline anisotropy is highly sensitive to Fe and Pt stoichiometry, the Fe/Pt sputtering power ratio was first systematically screened (table S1). In this design, the Fe target was fixed at 200 W and Pt target was operated at lower nominal power due to its higher sputtering yields under argon ion bombardment. Energy-dispersive spectroscopy (EDS) analysis showed that increasing Pt power from 120 to 180 W shifted the coating composition from Fe-rich to Pt-rich, whereas the Fe200-Pt140 condition yielded a near-equiatomic but slightly Fe-rich composition of approximately 55:45 Fe:Pt (Fig. S1). The nominal FePt film thickness was set to approximately 60 nm based on prior studies, which provides hard-magnetic actuation while limiting the remanence-driven aggregation observed for thicker FePt coatings (*17*). Step profilometry confirmed a deposited FePt thickness of 60.06 nm (Fig. S2). After annealing at 650 °C, the FePt coating underwent a clear structural transition from the disordered A1 phase to the chemically ordered L1_0_ phase (Fig. 2B). SEM revealed that the initially smooth as-deposited hemisphere became visibly granular after annealing, consistent with grain growth during ordering, while EDS confirmed Fe/Pt co-localization on the coated hemisphere (Fig. 2C). X-ray diffraction analysis (XRD) further verified phase ordering of all the Fe/Pt compositions. Fe200-Pt140 showed superlattice reflections at 2θ = 23.01° and 32.77°, assignable to the (100) and (110) peaks, respectively, and the split reflections at 46.98° (200) and 53.01° (002) were consistent with the tetragonal distortion of ordered Fe and Pt atoms (Fig. 2D and fig. S3). These diffraction features are characteristic signatures of chemically ordered L1_0_-FePt thin films reported in prior structural studies (*28*).

**Fig. 2.**
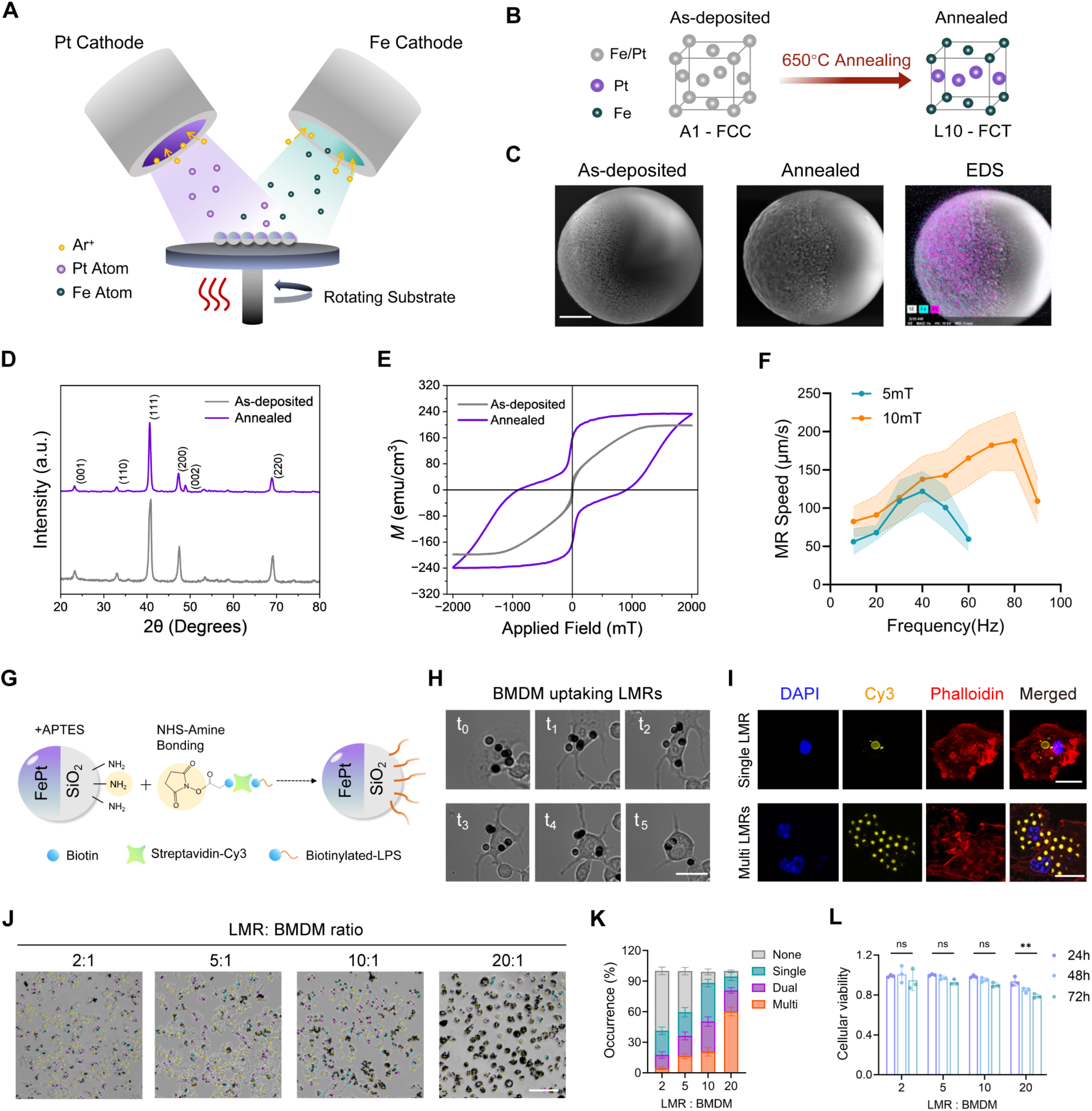
Fabrication and characterization of the Immunobot. (**A**) Schematic illustration of DC magnetron FePt co-sputtering strategy. (**B**) Lattice transformation from disordered A1 phase to ordered L1_0_ phase after annealing. (**C**) Morphology of L1_0_-FePt Janus particles before and after annealing at 650 °C. EDS scanning shows Fe (blue) and Pt (purple) atomic distribution on the film side. Scale bar, 1 µm. (**D**) XRD results showing the characteristic peaks variation between the FePt A1-phase and L1_0_-phase. (**E**) VSM characterization results of FePt coating before and after annealing showing the change from superparamagnetic state to ferromagnetic state. (**F**) The frequency sweep vs rolling velocity of FePt microrollers when actuated with 5 and 10 mT uniform rotating magnetic field showing the magnetic actuation of the fabricated microrollers. Data are mean ± SD (shaded areas). (**G**) Schematic illustration of biotin-streptavidin-based LPS conjugation to the non-magnetic side of microrollers, enabling selective biochemical modification. (**H**) Time-lapse brightfield microscopy images showing the phagocytosis of microrollers by BMDMs, with a time gap of 20 min between each figure. Scale bar, 20 µm. (**I**) Confocal microscopy images confirming the internalization of different numbers of LMRs by BMDMs. Scale bar, 20 μm. (**J**) Brightfield images showing BMDMs incubated with varying ratios of LMRs (2, 5, 10 and 20 folds) results in phagocytosis of single, dual and multi-roller immunobot generation. Panels show representative frames processed using a custom Python based recognition algorithm that identifies and labels immunobots according to their microroller count. (**K**) Statistical analysis of the occurrence of microroller number within a single cell after 24 h co-incubation. (**L**) Cell viability of BMDMs after co-incubation with varying ratio of microrollers for 24, 48, and 72 h. Data are mean ± SD, n = 3 biologically independent samples.

The structural ordering was accompanied by pronounced magnetic hardening. Vibrating sample magnetometer (VSM) measurements showed that, among all sputtering groups, Fe200-Pt140 produced the strongest remanence and the largest coercivity, reaching Mr = 169.11 emu/cm³ and Hc = 717.90 mT. Compared with its as-deposited state, the annealed coating also exhibited markedly increased hysteresis squareness (Fig. 2E). With further increasing Fe content, saturation magnetization continued to rise, whereas both hysteresis squareness and coercivity declined, indicating that Fe enrichment did not further improve effective magnetic hardness. Previous research has also shown that stable magnetic actuation is better correlated with high coercivity and remanence than with saturation magnetization alone (fig. S4). After characterization of the structural and material properties, magnetic actuation of the fabricated microrollers was tested with a custom-made five-coil electromagnetic system (fig. S5). Our MRs showed a frequency-dependent rolling motion over a broad operating window (movie S1). To evaluate locomotion performance under conditions relevant to clinical translation, where large-workspace electromagnetic systems must limit dynamic field strengths to minimize Joule heating and tissue eddy currents, a biologically safe field of 10 mT was selected as the benchmark (*29*, *30*). At this clinically practical field strength, the translational speed of the MRs increased with driving frequency, reaching a maximum average value of 187.70 µm/s, while this robust synchronous rolling was maintained up to a step-out frequency of 80 Hz (Fig. 2F).

To enable the magnetic microrollers to act as a biochemical regulator of macrophage phenotype and functionality, the non-magnetic SiO₂ hemisphere of the Janus microroller was conjugated with LPS through sequential amine coupling and biotin-streptavidin interaction to form the LPS-modified FePt microroller (LMR) (Fig. 2G). The Cy3-labeled streptavidin enabled indirect visualization of the LPS distribution on the microroller surface. The successful stepwise fabrication of the engineered microparticles was first validated through zeta potential measurements. Zeta potential analysis showed a progressive surface charge change from SiO₂ microspheres to FePt coated MRs and LMRs, with LMRs displaying the most negative potential (−18.2 mV), consistent with the introduction of abundant negatively charged phosphate and carboxyl groups on the particle surface (fig. S6A). Inductively coupled plasma mass spectrometry ICP-MS further confirmed tunable biotin-LPS immobilization on the microrollers, with the loaded amount increasing with input concentration and reaching ∼320 μg per 10⁷ MRs at the highest tested condition (fig. S6B). This value was therefore defined as the practical upper loading level achieved under the current coupling protocol. Notably, the surface conjugated LPS should be distinguished from freely soluble LPS, the quantified value was considered a nominal particle associated payload.

After preparation of the LMRs, BMDMs were isolated from the femurs of mice (fig. S7) and co-incubated with LMRs for 24 hours to generate immunobots. Time-lapse imaging captured progressive engulfment of micron sized LMRs after cell contact (Fig. 2H, movie S2), while fluorescence imaging confirmed intracellular localization of both single and multiple payloads (Fig. 2I). We hypothesized that excessive intracellular LMR loading may impair BMDM viability. To quantify and control LMR uptake, BMDMs were incubated with LMRs at input ratios ranging from 2:1 to 20:1. Based on the number of internalized LMR per cell, the resulting BMDMs were classified as single-, dual-, and multi-loaded immunobots, corresponding to cells containing one, two, and three or more LMRs, respectively. Increasing the input ratio progressively reduced the fraction of unloaded cells while increasing the frequencies of multi-payload cells, causing a markedly heterogeneous distribution of loading states within the final immunobot population (Fig. 2 J and K). Under the conditions of 2:1, 5:1 and 10:1 loading, BMDMs tolerated intracellular loading well, with viabilities remaining above 85% from 24 to 72 h. In contrast, the highest loading condition (20 LMRs per BMDM) showed a clear late decline, with only 77.0 ± 4.12% of the cells remaining viable at 72 h (Fig. 2L), indicating that excessive payload imposed a measurable cytological burden. To balance cell viability with intracellular loading efficiency at a population level, we selected 10 LMRs per BMDM as the input ratio for subsequent experiments. At this ratio, 37.78% ± 3.70% and 29.75% ± 4.60% of BMDMs were converted into single- and dual-roller immunobots, respectively.

### LMR payload dictates immunobot propulsion, penetration, and cellular migration dynamics

Having established immunobot formation and identified an optimal LMR input ratio, we next examined how payload number influenced magnetic actuation at a cellular level. Considering L1_0_-FePt is a hard-magnetic material with strong remanent magnetization, we assumed the torque generated under a rotating magnetic field is governed by the effective vector sum of intracellular magnetic moments, τ_effective_ = (∑_i_ m_i_) × B, rather than by the total magnetic payload alone. After phagocytosis, randomly oriented and lysosomally clustered LMRs may undergo partial vector cancellation, while close-packed microroller assemblies may further reduce the externally available magnetic moment through dipole–dipole interactions and flux closure configurations. We therefore introduced a pulsed re-magnetization step after cellular uptake to realign the intracellular magnetic moments and increase the effective net moment of the immunobots (Fig. 3A). Before re-magnetization, increasing the number of internalized LMRs did not produce a proportional improvement in propulsion. Single-, dual-, and multi-roller immunobots displayed comparable step-out frequencies of approximately 20 Hz. Single-roller immunobots reached 72.84 ± 11.14 μm/s at this step-out frequency, whereas the maximum velocities of dual- and multi-roller immunobots were 108.55 ± 14.93 μm/s and 131.43 ± 25.21 μm/s, corresponding to only approximately 1.48-fold and 1.82-fold increases relative to single-loaded cells, respectively. Following re-magnetization, dual- and multi-payload immunobots showed markedly improved magnetic response, with step-out frequencies increasing from 20 to 35 Hz. Their maximum rolling speeds also increased to 132.58 ± 19.31 μm/s and 206.64 ± 30.97 μm/s, respectively, with the strongest enhancement observed in multi-payload immunobots (Fig. 3, B and C; movie S3). The re-magnetized immunobots retained controllable motion across biologically relevant substrates. On glass, they followed prescribed trajectories with stable rolling motion under a 10 Hz, 10 mT rotating magnetic field (Fig. 3D). Similar control was maintained on HUVEC monolayers and in whole blood, although the rolling speed decreased under cellular interfacial resistance and blood-associated fluidic resistance (Fig. 3, E and F; movie S4). As prolonged magnetic actuation and contact with complex biological interfaces could potentially influence macrophage viability and activation state, we next examined whether the operating condition used here altered BMDM status. Importantly, continuous actuation below the step-out frequency did not cause obvious BMDM viability loss or phenotype shifting (fig. S8).

**Fig. 3.**
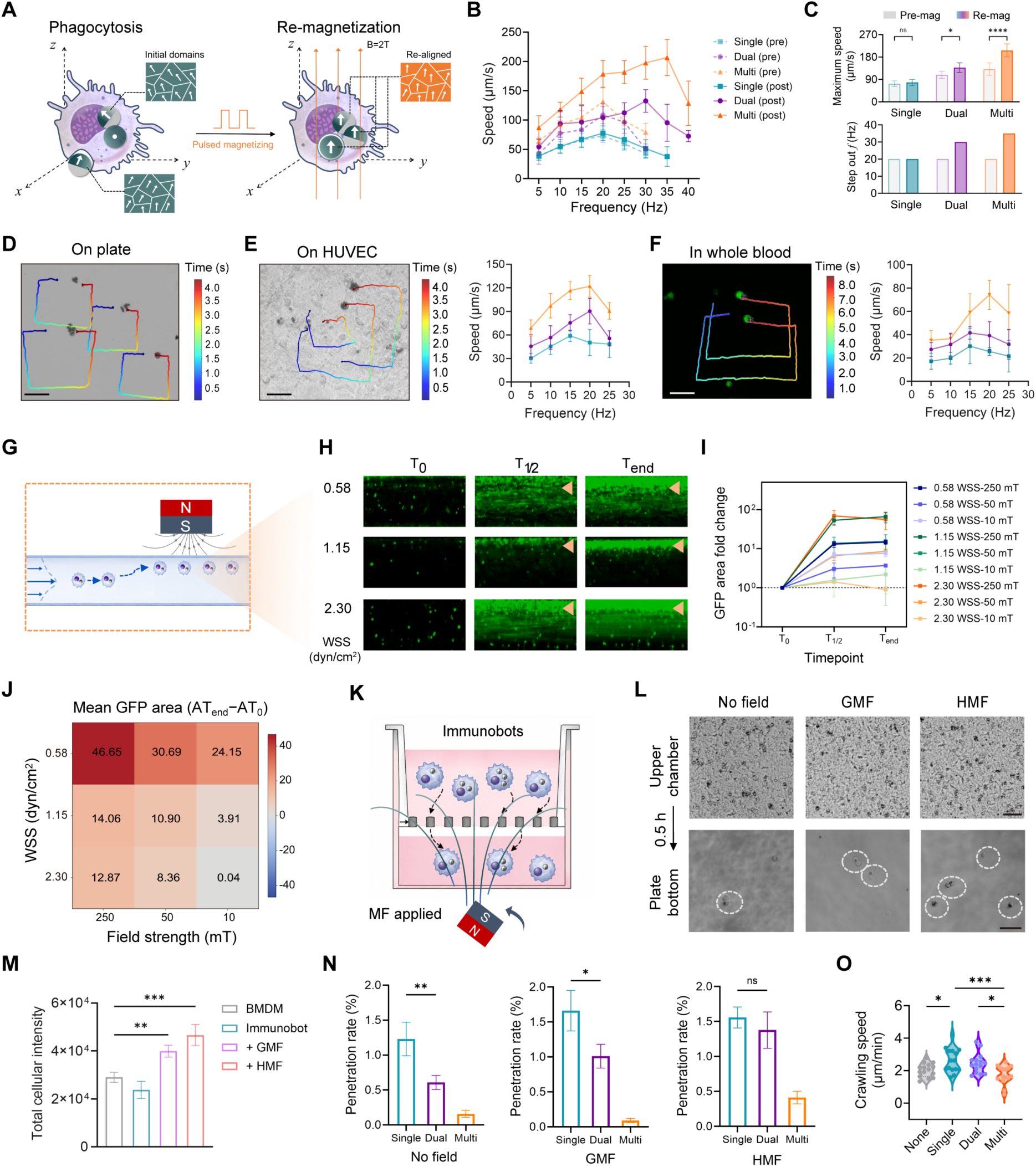
Payload-dependent magnetic actuation, barrier penetration, and basal motility of immunobots. (**A**) Schematic illustration of intracellular dipole realignment by post-engulfment pulsed remagnetization of internalized LMRs. (**B**) Frequency-dependent rolling speed of pre-magnetized and remagnetized single-, dual-, and multi-payload immunobots under externally applied 10 mT uniform magnetic field. Data are mean ± SD (shaded areas), n*=* 20. (**C**) Step-out frequency and maximum rolling speed of immunobots before and after remagnetization, showing the effect of remagnetization on increasing the resulting rolling speed. (**D** to **F**) Representative trajectories and speed-frequency profiles of remagnetized immunobots on glass, on HUVEC monolayers, and in whole blood. Data are mean ± SD (shaded areas), n=10. Scale bar, 50 µm. (**G**) Schematic illustration of the *in vitro* microfluidic assay. Immunobots were loaded into a straight microchannel, while an external permanent magnet positioned adjacent to the channel generates a magnetic field gradient that biases cells toward the magnet-facing wall and promotes local retention against flow. Blue arrows indicate the flow rate profile within the channel. (**H**) Representative fluorescence images acquired from the microchannel at wall shear stress of 0.58, 1.15 and 2.3 dyn/cm^2^ under 250 mT field at the T_0_, T_1/2,_ and T_end_. Orange markers indicate the magnet-facing region. Scale bar, 300 μm. (**I**) Quantification of retained fluorescence area fold change normalized to the corresponding T_0_ signal. (**J**) Heatmap of normalized retention fluorescence area across the field–WSS matrix. (**K**) Schematic of the 8 µm-pore transwell assay for the analysis of immunobot penetration. (**L**) Representative microscopy images showing immunobots in the upper chamber and transmigrated immunobots in the lower chamber in the presence and absence of a magnetic field. Scale bar, 50 μm. (**M**) Total fluorescence intensity of transmigrated immunobots detected in the lower chamber. Data are mean ± SD, *n* = 3 replicates. (**N**) Normalized penetration rate under no field, static field, and hybrid field conditions demonstrating the significantly reduced penetration abilities of multiroller-loaded immunobots. (**O**) Comparison of basal crawling speeds of unloaded BMDMs and immunobots carrying different numbers of internalized microrollers, showing the multiroller-loaded immunobots with compromised migration abilities. Statistical significance was determined by one-way ANOVA with Tukey’s multiple comparisons test. \**P* < 0.05; \*\**P* < 0.01; \*\*\**P* < 0.001; ns, not significant.

To determine whether magnetically responsive immunobots could be captured from flow before barrier interaction, we established a vessel-mimetic cylindrical microchannel assay using a 300 μm diameter PDMS channel perfused with serum containing carboxyfluorescein succinimidyl ester (CFSE) stained immunobot suspensions (Fig. 3G). The gradient magnetic field was generated by placing a permanent magnet adjacent to the chamber, producing a spatially varying magnetic flux density that directed immunobots toward the high-field region. To approximate low shear vascular conditions relevant to tumor-associated microcirculation (*31–33*), the assay was performed at linear flow velocities of 1.8, 3.6, and 7.2 mm/s, corresponding to estimated wall shear stresses (WSS) of 0.58, 1.15, and 2.3 dyn/cm², respectively. Time-lapse fluorescence imaginga revealed progressive immunobot enrichment along the magnet-facing side of the channel during perfusion (Fig. 3H, figs. S9 and 10). The accumulation pattern depended jointly on hydrodynamic load and magnetic field strength. Quantification of regions of interest (ROI) area showed a time-dependent fluorescence signal increase under most field–flow conditions, but the magnitude of accumulation did not scale linearly with WSS (Fig. 3I and movie S5). This non-uniform response is consistent with the competing effects of cell delivery within the flow and the shear induced washout. Increasing flow velocity increases the number of immunobots delivered to the magnetic capture region per unit time, while simultaneously reducing the residence time and capture probability of individual cells. Under low WSS, the prolonged residence time permitted detectable retention even at 10 mT. At intermediate WSS, washout became more pronounced, whereas the increased cell influx was insufficient to fully compensate for reduced capture probability. At the highest WSS, weak field retention remained near the threshold, but residual signal could still be detected, likely reflecting increased convective delivery into the ROI rather than efficient stable capture. The 3 × 3 field-WSS matrix, therefore, served as an operational map of immunobot retention, identifying low-field/high-shear conditions as near-threshold regimes and showing that increasing magnetic field strength partially restored retention under elevated hydrodynamic load (Fig. 3J). Together, these results indicate that immunobot sequestration under flow is governed by a combination of magnetic attraction, delivery amount, residence time, and flow-induced re-enrichment. This operational map may further inform *in vivo* magnetic guidance, where immunobot entry into the target vasculature is likely to be limited and temporally variable, suggesting that repeated local attraction may improve arrival and vessel wall retention.

However, such vascular capture represents only the first step toward tissue delivery, we next examined whether magnetic actuation could further promote barrier crossing after capturing. Transmigration was evaluated using an 8 µm-pore transwell assay under no field, gradient magnetic field (GMF), or hybrid magnetic field (HMF) consisting of a gradient plus rotating field (Fig. 3, K and L). To account for differences in the initial occurrence of single-, dual-, and multi-payload immunobots in the upper chamber, penetration rates were calculated by normalizing the number of transmigrated cells in each payload category to its estimated input number before field application. Compared with unloaded BMDMs, immunobots in the no-field condition exhibited lower transmigration, consistent with the mechanical burden imposed by rigid intracellular LMR payloads during confined pore crossing (Fig. 3M). Magnetic stimulation partially overcame this limitation in a field-mode-dependent manner, with HMF producing the highest total fluorescence signal and microscopy-normalized penetration across groups (Fig. 3M).

Under HMF conditions, single-, dual-, and multi-payload immunobots reached improved penetration rates of approximately 1.56%, 1.37%, and 0.47%, respectively, compared with 1.23%, 0.61%, and 0.16% in the corresponding no-field controls (Fig. 3N). Thus, even multi-payload immunobots, which were otherwise limited by their higher intracellular cargo burden, showed improved barrier crossing under hybrid actuation. To further determine whether payload-dependent penetration differences were associated with altered intrinsic motility, we tracked basal crawling in the absence of magnetic actuation. Relative to unloaded macrophages (2.04 ± 0.42 µm/min), single- and dual-payload immunobots showed slightly higher average crawling speeds (2.63 ± 0.74 and 2.45 ± 0.61 µm/min, respectively), whereas the multi-payload group decreased to 1.75 ± 0.56 µm/min (Fig. 3O). This trend raises the possibility that limited LMR uptake may modestly prime macrophage motility, rather than acting solely as a passive mechanical load. The modest increase in crawling speed observed in single- and dual-loaded cells further suggests that limited LMR uptake may weakly improve macrophage motility, potentially through particle-induced adhesion remodeling or LPS-associated actin reorganization (*34*, *35*).

### LMR-driven iron metabolism mediates M1-like immunobot polarization

Given the well-established immunostimulatory activity of LPS and the emerging role of iron-associated signals in driving proinflammatory macrophage programming, we next investigated whether the incorporation of LPS-modified FePt microrollers endows immunobots with an enhanced M1-like polarization phenotype (*18*, *36*). Flow cytometry and corresponding mean fluorescence intensity (MFI) analysis showed that immunobots significantly upregulated the M1-associated marker CD86 relative to baseline macrophage colony-stimulating factor (M-CSF)-differentiated BMDMs, with a greater induction effect than that achieved by direct stimulation with free LPS. In contrast to the control group, immunobots did not induce an increase in the M2 marker CD206 (Fig. 4, A and B and fig. S11A). Cytokine secretion profiles further supported this pro-inflammatory phenotypic shift. Relative to the control, immunobots significantly promoted the secretion of tumor necrosis factor-α (TNF-α), interleukin-6 (IL-6), and interleukin-12 (IL-12). Notably, the IL-6 level was markedly higher than that in the LPS-only treatment group (Fig. 4C), whereas the anti-inflammatory cytokine interleukin-10 (IL-10) was maintained at a basal low level (Fig. 4D). Importantly, time-course monitoring at 24 h, 48 h, and 72 h revealed highly consistent trends in both flow cytometry and enzyme-linked immunosorbent assay (ELISA) results (figs. S11 and S12), indicating that this internalization process successfully endowed immunobots with a potent and stable M1 polarization phenotype.

**Fig. 4.**
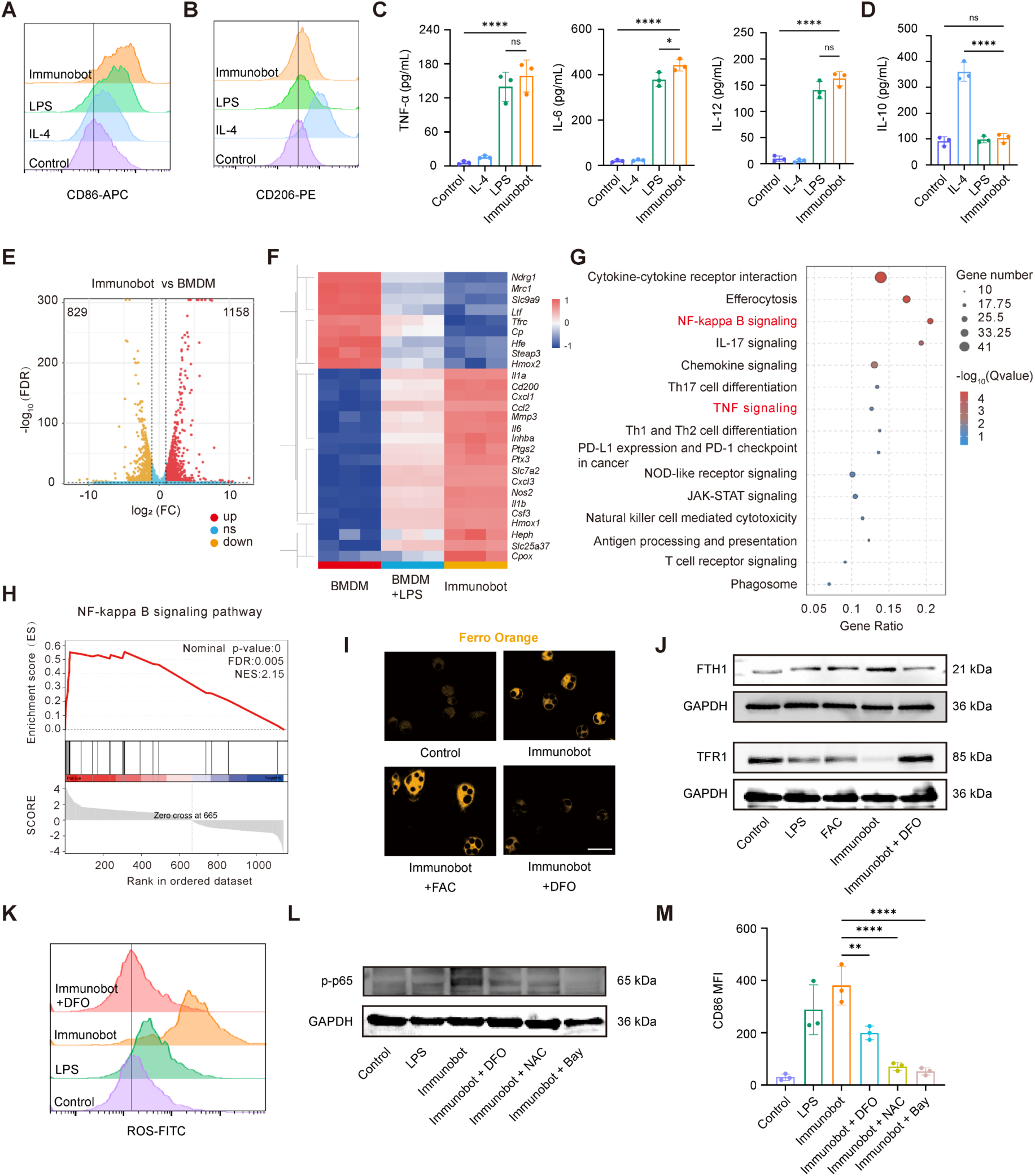
Immunobots acquire an iron-dependent M1-like phenotype. (**A** and **B**) Representative flow cytometry histograms showing the expression of CD86 (A) and CD206 (B) in BMDMs at 24 h following the indicated treatments. Control, M-CSF-differentiated BMDMs without additional IL-4 or LPS stimulation and without LMR loading. (**C** and **D**) Quantification of the proinflammatory cytokines TNF-α, IL-6, and IL-12 (C) and the anti-inflammatory cytokine IL-10 (D) in culture supernatants at 24h following the indicated treatments. (**E**) Volcano plot showing differentially expressed genes (DEGs) between BMDMs and immunobots. Red and yellow dots indicate upregulated and downregulated genes, respectively. (**F**) Hierarchical clustering heatmap of representative M1 polarization and iron metabolism-related genes in BMDMs, LPS-treated BMDMs, and immunobots. **(G)** KEGG pathway enrichment analysis of DEGs between immunobots and baseline BMDMs. Dot size indicates the number of genes, and color indicates −log₁₀ (*q* value). **(H)** GSEA showing enrichment of the NF-κB signaling pathway in immunobots relative to baseline M-CSF-differentiated BMDMs. (**I**) Representative Ferro Orange staining images showing intracellular labile Fe²⁺ levels in BMDMs after the indicated treatments. Scale bar, 20 μm. (**J**) Western blot analysis of iron metabolism-related proteins FTH1 and TFR1 in BMDMs at 24 h following the indicated treatments. GAPDH was used as a loading control. (**K**) Representative flow cytometry histograms showing intracellular ROS levels detected by DCFH-DA staining after the indicated treatments. (**L**) Western blot analysis of p65 phosphorylation at Ser536 in BMDMs in the indicated groups. DFO, NAC, and BAY 11-7082 were used to inhibit iron accumulation, ROS, and NF-κB signaling, respectively. GAPDH was used as a loading control. (**M**) Quantification of CD86 MFI in BMDMs after the indicated treatments. Data are presented as mean ± SD , n = 3 biologically independent samples. Statistical significance was determined by one-way ANOVA with Tukey’s multiple comparisons test. \**P* < 0.05, \*\**P* < 0.01, \*\*\*\**P* < 0.0001; ns, not significant.

To further define the molecular basis of immunobot activation, we performed bulk RNA-seq analysis of BMDMs, LPS-treated BMDMs, and immunobots. Venn, differential expression, and GO enrichment analyses revealed that immunobots largely recapitulated the LPS-induced inflammatory transcriptional program while acquiring additional LMR-associated transcriptional features related to iron ion transport and iron homeostatic regulation (Fig. 4E and fig. S13A to C). Specifically, 1158 genes were upregulated, and 829 genes were downregulated in the immunobot group compared to BMDM. Hierarchical clustering showed that immunobots markedly upregulated canonical M1 inflammatory genes, including *Nos2*, *Il1b*, and *Il6*, while downregulating the M2 marker *Mrc1* (Fig. 4F), suggesting that both LPS stimulation and LMR loading contribute to the conversion of macrophages toward a proinflammatory M1-like state. Notably, compared with both baseline BMDMs and LPS-stimulated BMDMs, immunobots exhibited a coordinated yet nonuniform transcriptional response in iron homeostasis-related genes, characterized by decreased expression of iron uptake-associated genes, including *Tfrc* and *Ltf*, and increased expression of stress-response and intracellular iron-handling genes, including *Hmox1*, *Cp*, *Slc25a37*, and *Cpox*. This pattern suggests that intracellular processing of FePt particles may increase the labile iron pool and induce compensatory iron buffering, redox adaptation, and mitochondrial iron redistribution. Further Kyoto Encyclopedia of Genes and Genomes (KEGG) pathway enrichment analysis and gene set enrichment analysis (GSEA) revealed significant enrichment of the nuclear factor kappa B (NF-κB) signaling pathway, along with activation of other inflammation-related pathways, including TNF, IL-17, chemokine signaling, and cytokine-cytokine receptor interaction pathways (Fig. 4G, H). Together, these transcriptomic signatures suggest that immunobot formation couples inflammatory activation with iron homeostatic remodeling, potentially allowing LPS priming and LMR-associated intracellular responses to reinforce macrophage polarization through NF-κB signaling.

Guided by transcriptomic evidence of altered iron metabolism, we next examined intracellular iron homeostasis in immunobots. Ferro Orange staining revealed an increased intracellular labile Fe²⁺ pool, which was further elevated by exogenous iron loading with ferric ammonium citrate (FAC) and reduced by iron chelation with deferoxamine (DFO) (Fig. 4I and fig. S14A). To test whether this increase could arise from LMR degradation, we performed an *in vitro* assay under phagolysosome-mimicking acidic conditions. The residual FePt coating showed decreased Fe content, while ICP-MS detected increased Fe release in the supernatant, without a significant loss of magnetic responsiveness (fig. S15). These results suggest that LMR degradation contributes to intracellular labile iron accumulation while preserving magnetic functionality. Protein-level validation further delineated the nature of this metabolic rewiring. Immunobots significantly upregulated ferritin heavy chain (FTH1), which is responsible for iron storage, and correspondingly downregulated transferrin receptor 1 (TFR1), which mediates extracellular iron uptake. Treatment with DFO successfully reversed this expression profile (Fig. 4J and fig. S14B). This high-storage, low-uptake iron-handling pattern is consistent with the iron-retentive phenotype of proinflammatory M1-like macrophages, which may facilitate the accommodation of iron-associated stress and contribute to ferroptosis resistance while preserving inflammatory function (*19*, *20*, *37*). Given the increased intracellular labile Fe²⁺ pool in immunobots, we next examined whether iron-associated redox activity contributed to oxidative stress. Flow cytometric analysis revealed markedly elevated ROS levels in immunobots, which were significantly reduced by DFO treatment, supporting a substantial contribution of labile iron to ROS accumulation, likely through Fe²⁺-mediated Fenton chemistry (Fig. 4K and fig. S14C). To explicitly determine whether iron metabolic remodeling and the ROS burst cascade are a prerequisite for triggering NF-κB-mediated M1 polarization, we conducted targeted intervention experiments. Western blot analysis showed increased phosphorylation of p65 in immunobots, suggesting activation of NF-κB signaling. Notably, restricting iron loading with DFO, neutralizing downstream ROS with N-acetylcysteine (NAC), or inhibiting NF-κB with BAY 11-7082 each substantially attenuated this phosphorylation signal (Fig. 4L and fig. S14D). More importantly, quantitative flow cytometry analysis verified that all three intervention strategies effectively blocked the immunobot-driven M1 polarization trajectory, erasing its characteristic high CD86 expression (Fig. 4M and fig. S14E). Together, these results indicate that the potent M1-like polarization of immunobots is driven by the synergistic effects of LMR-delivered LPS-mediated inflammatory stimulation and Fe²⁺-associated redox stress, which collectively enhance ROS accumulation and NF-κB signaling.

### Immunobots induce ferroptosis-associated immunogenic cell death in tumor cells

Having demonstrated that immunobots acquire a proinflammatory M1-like phenotype accompanied by enhanced iron-associated redox activity, we next examined whether these activated cells could transmit cytotoxic signals to tumor cells independently of direct cell-cell contact. A noncontact coculture system was established, with baseline M-CSF-differentiated BMDMs (M0), LPS-stimulated BMDMs (M1), or immunobots in the upper chamber and MC38 tumor cells in the lower chamber (Fig. 5A). Live/dead staining showed that immunobots induced more pronounced MC38 cell death than M0 or conventional M1 macrophages (Fig. 5B and fig. S16A). Consistently, Annexin V/PI flow cytometry revealed a marked increase in late apoptotic/secondary necrotic MC38 cells after immunobot coculture, with the Annexin V⁺PI⁺ population increasing from 11.2% in the M1 group to 29.0% in the immunobot group (Fig. 5C). Because immunobots carry LPS-modified FePt microrollers and undergo iron metabolic remodeling during their generation, we next asked whether their antitumor effect involved iron-related tumor cell death. The increase in MC38 cell death was partially attenuated by ferrostatin-1 (Fer-1), a lipid radical-trapping inhibitor of ferroptosis, suggesting that immunobot-mediated tumor cell killing involves a ferroptosis-related, lipid peroxidation-dependent component.

**Fig. 5.**
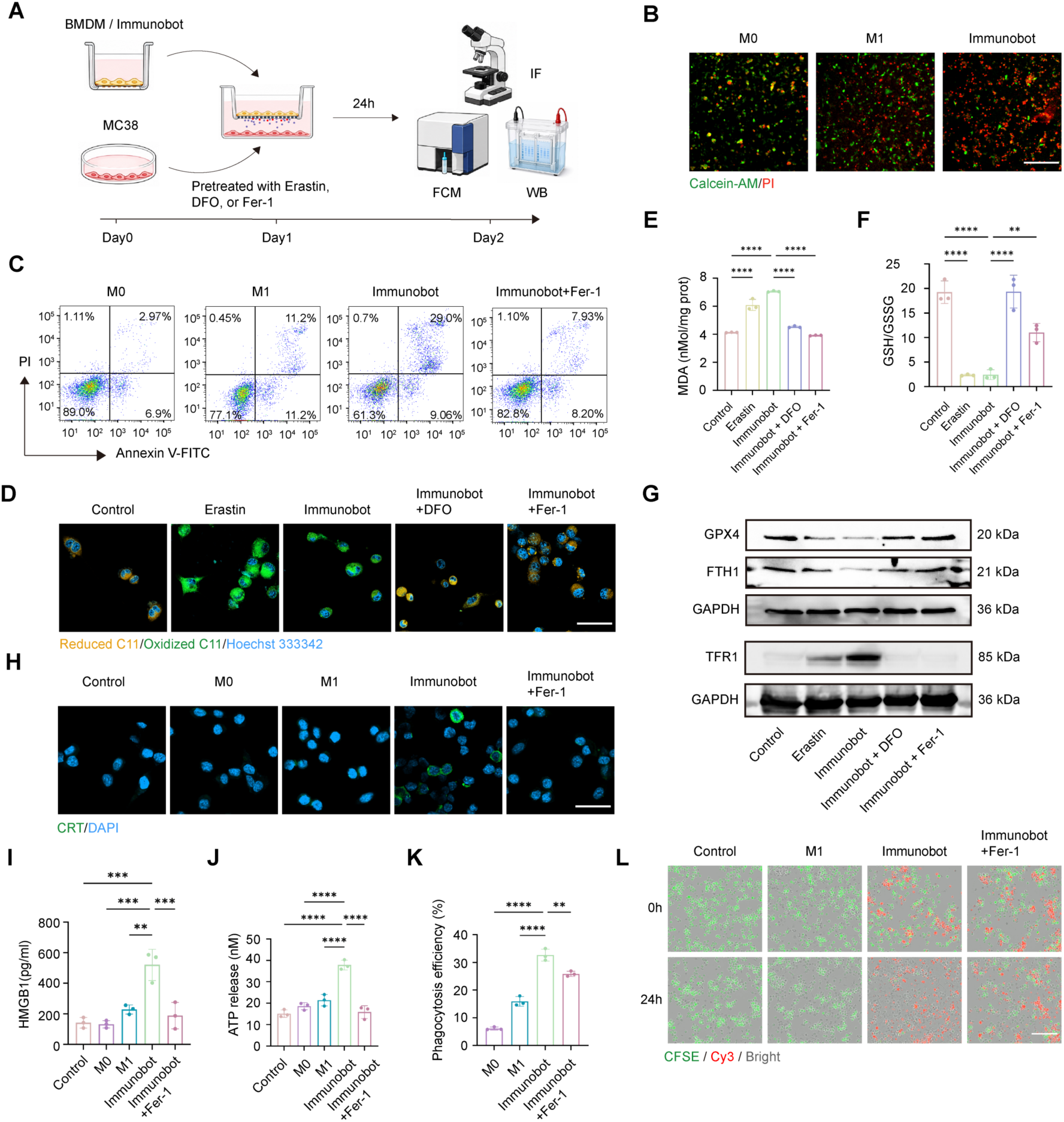
Immunobots induce ferroptosis-associated immunogenic cell death in tumor cells. (**A**) Schematic of the non-contact coculture system used to evaluate immunobot-mediated tumor cell killing. BMDMs or immunobots were seeded in the upper chamber, and MC38 tumor cells were cultured in the lower chamber on day 0. Where indicated, MC38 cells were pretreated with erastin, DFO, or Fer-1 on day 1 before coculture, followed by immunofluorescence imaging, flow cytometry, or western blot. (**B**) Live/dead staining of MC38 cells after coculture with M0 macrophages, M1 macrophages, or immunobots. Live cells were labeled with calcein-AM, and dead cells were labeled with PI. Scale bar, 200 μm. **(C)** Representative Annexin V-FITC/PI flow cytometry plots showing MC38 cell death after coculture with the indicated macrophage groups, with or without Fer-1. (**D**) Representative C11-BODIPY staining images showing lipid ROS accumulation in MC38 cells. Untreated MC38 cells served as the control, and erastin was used as a ferroptosis-positive control. Scale bar, 50 μm. (**E** and **F**) Quantification of MDA levels (E) and GSH/GSSG ratios (F) in MC38 cells after the indicated treatments. (**G**) Western blot analysis of ferroptosis- and iron metabolism-related proteins in MC38 cells, including GPX4, FTH1, and TFR1. GAPDH was used as a loading control. (**H**) Immunofluorescence staining of CRT exposure on MC38 cells following the indicated coculture treatments. Nuclei were stained with DAPI. Scale bar, 50 μm. (**I** and **J**) Quantification of HMGB1 (I) and ATP (J) release from MC38 cells after the indicated coculture treatments. (**K**) Representative flow cytometry plots comparing the phagocytosis of MC38 cells by the indicated groups. (**L**) Incucyte-based live-cell imaging of CFSE-labeled MC38 cells during coculture. Cy3 signal indicates LMRs. Scale bar, 100 μm. Data are presented as mean ± SD, n = 3 biologically independent samples. Statistical significance was determined by one-way ANOVA with Tukey’s multiple comparisons test. \*\**P* < 0.01; \*\*\*\**P* < 0.0001.

Because ferroptosis is characterized by iron-dependent lipid peroxidation rather than by loss of cell viability alone (*23*, *38*, *39*), we next examined lipid peroxidation and ferroptosis-associated molecular changes in MC38 cells. ROS-FITC flow cytometry showed that immunobot coculture increased intracellular ROS levels in MC38 cells (fig. S16B). C11-BODIPY staining further showed that immunobot treatment increased lipid ROS accumulation in MC38 cells, supporting the induction of lipid peroxidation (Fig. 5D and fig. S16C). This lipid peroxidation phenotype was further supported by increased malondialdehyde (MDA), a terminal product of lipid peroxidation, and by a decreased GSH/GSSG ratio, indicating impaired antioxidant buffering capacity (Fig. 5E and F). Both the iron chelator DFO and Fer-1 attenuated these changes, supporting an iron- and lipid peroxidation-dependent process. Western blot analysis further substantiated the ferroptotic phenotype. Immunobot treatment significantly decreased GPX4 expression by 5-fold, indicating impaired enzymatic detoxification of lipid peroxides. In parallel, immunobots reshaped iron-homeostasis proteins in MC38 cells, as evidenced by decreased FTH1 and increased TFR1 expression (Fig. 5G and fig. S16D). This pattern suggests disrupted iron homeostasis, characterized by reduced iron-storage capacity and enhanced iron-uptake potential, thereby favoring iron-dependent lipid peroxidation and ferroptosis susceptibility. DFO or Fer-1 treatment partially reversed these protein changes, further supporting that immunobot-induced MC38 cell death is associated with iron-dependent lipid peroxidation.

We next examined whether the ferroptosis-associated death of MC38 cells was accompanied by immunogenic danger signaling. Because the immunogenic potential of dying tumor cells is associated with the coordinated emission of damage-associated molecular patterns (DAMPs), we assessed surface-exposed calreticulin (CRT), extracellular ATP, and HMGB1 release as canonical ICD-associated markers (*40–42*). Immunofluorescence staining showed that immunobot coculture markedly promoted CRT exposure on MC38 cells, whereas Fer-1 reduced this effect (Fig. 5H and fig. S16E). In parallel, immunobot treatment increased HMGB1 and ATP release into the culture supernatant, and both responses were attenuated by Fer-1 treatment (Fig. 5I and J). These results indicate that immunobot-induced ferroptotic stress is coupled with ICD-associated DAMP emission.

Because surface-exposed CRT functions as a pro-phagocytic “eat-me” signal that facilitates tumor cell recognition and engulfment by macrophages, we further examined whether immunobots exhibited enhanced phagocytosis of MC38 tumor cells in a direct coculture setting (*43*, *44*). To assess tumor cell phagocytosis, Cy3-labeled immunobots were directly cocultured with CFSE-labeled MC38 cells. Flow cytometry analysis showed that immunobots increased the proportion of Cy3⁺CFSE⁺ phagocytic cells from 15.90% in the M1 macrophage group to 32.63%, whereas Fer-1 treatment partially attenuated this effect, reducing the proportion to 25.87% (Fig. 5K and fig. S17). These results suggest that lipid peroxidation-dependent ferroptotic processes contribute to immunobot-enhanced tumor cell phagocytosis. Finally, Incucyte live-cell imaging further showed a time-dependent reduction of CFSE⁺ MC38 cells after immunobot coculture, whereas Fer-1 partially preserved tumor cell survival and growth (Fig. 5L). Together, these findings indicate that immunobots promote tumor cell elimination through ferroptosis-linked immunogenic remodeling, marked by soluble cue-induced lipid peroxidation, ICD-related DAMP emission, and Fer-1-sensitive macrophage phagocytosis.

### Hybrid magnetic actuation enhances immunobot delivery and antitumor efficacy *in vivo*

To evaluate the tumor-targeted delivery capability and therapeutic efficacy of immunobots *in vivo*, we established a subcutaneous syngeneic colorectal tumor model in C57BL/6 mice. When tumors reached approximately 100 mm³ on day 7 after tumor inoculation, mice were randomly assigned to six groups: G1-Saline, G2-M1 macrophages, G3-LMR-Immunobot, G4-MR-Immunobot + HMF, G5-LMR-Immunobot + GMF, and G6-LMR-Immunobot + HMF. Saline, M1 macrophages, or the indicated immunobot formulations were administered via tail-vein injection. Where indicated, mice were subsequently subjected to either a gradient magnetic field alone or a hybrid magnetic field combining static and rotating magnetic fields (movie S6), the latter of which was shown to effectively enhance the penetration ability of immunobots *in vitro* (Fig. 3M, Fig. 6A, B and fig. S18). Longitudinal *in vivo* imaging system (IVIS) was used to assess the biodistribution of the DiD-labeled BMDMs and immunobots (movie S7). Fluorescence images showed detectable fluorescence signals in the tumor region of all groups after administration, which gradually decreased after day 1 (Fig. 6C, D). Among all groups, the LMR-Immunobot + HMF group (G6) showed the strongest tumor-localized fluorescence throughout the observation period, indicating enhanced tumor recruitment and local retention under hybrid magnetic actuation. *Ex vivo* IVIS imaging at the study endpoint further revealed that fluorescence signals were mainly distributed in the liver and spleen, with no evident signal detected in the heart, lungs, kidneys, or whole excised tumors (fig. S19), suggesting predominant uptake and clearance through the reticuloendothelial system without widespread nonspecific accumulation in peripheral organs. Consistent with the *in vivo* imaging results, the G6 group showed the most pronounced antitumor activity. Tumor growth curves demonstrated that tumor progression was most effectively suppressed in G6, which also exhibited the smallest tumor volume and the lowest tumor weight at the endpoint (Fig. 6E to G and fig. S20). To further assess intratumoral delivery efficiency, FePt accumulation in tumor tissues was quantified and examined histologically. ICP-MS analysis showed that all magnetically guided groups exhibited high intratumoral Pt content, with G6 showing the highest level (Fig. 6H). Similarly, G6 also showed stronger Prussian blue staining (Fig. 6I) and more persistent DiD-associated fluorescence within the tumor tissue (Fig. 6J). The CD31/DiD images revealed that the DiD signal was localized not only to CD31-positive vascular regions but also to extravascular tumor parenchyma, indicating that under hybrid magnetic actuation, immunobots could not only efficiently access tumor-associated vasculature but also further penetrate into the tumor parenchyma (Fig. 6J). In addition, CD31 staining and quantitative analysis showed a marked reduction in tumor vascular area in G6, suggesting substantial vascular damage induced by this treatment (Fig. 6J and K). Together, these results demonstrate that hybrid magnetic actuation markedly improves the intratumoral deposition, retention, and tissue penetration of immunobots.

**Fig. 6.**
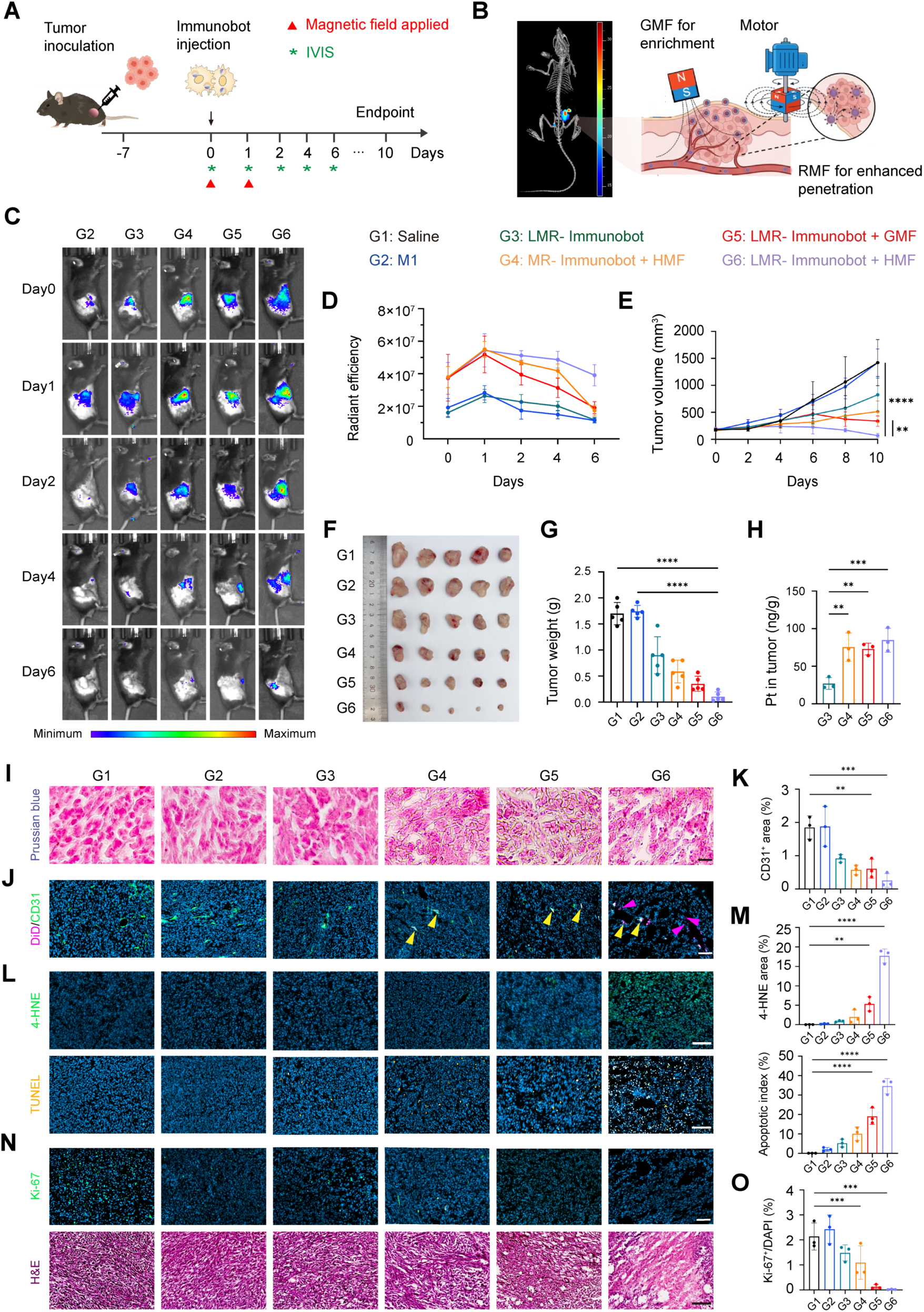
Hybrid magnetic actuation enhances the tumor targeting and antitumor efficacy of immunobots in a murine colorectal cancer model. (**A**) Schematic timeline of the animal experiment in MC38 subcutaneous tumor-bearing mice. (**B**) Schematic illustration of subcutaneous tumor localization and the application of the gradient magnetic field (GMF) and rotating magnetic field (RMF) with a representative microCT-IVIS co-registered image of tumor-bearing mice. (**C**) Representative IVIS images acquired at Day 0 and Day 1 after each of the two magnetic field applications, as well as at Days 2, 4, and 6. (**D**) Quantification of DiD fluorescence in the tumor ROI as average radiant efficiency, measured at the indicated time points [(p/s/cm²/sr)/(μW/cm²)]. (**E**) Tumor growth curves showing tumor volume measured on Days 0, 1, 2, 4, and 6, normalized to the tumor volume at Day 0. (**F**) Photographs of excised tumors collected at the study endpoint. (**G**) Tumor weights measured at the endpoint. (**H**) Pt content in tumors measured by ICP-MS. (**I**) Representative Prussian blue staining of tumor sections showing intratumoral ferric iron deposition. Scale bar, 20 μm. **(J)** Representative fluorescence images of tumor sections stained for CD31 together with the DiD signal. Yellow arrowheads indicate DiD signals overlapping with CD31-positive blood vessels, whereas pink arrowheads indicate DiD signals located within the tumor parenchyma. Scale bar, 100 μm. **(K)** Quantification of the CD31-positive area fraction in tumor sections. **(L)** Representative images of 4-HNE staining (top) and TUNEL staining (bottom) in tumor sections. Scale bar, 100 μm. **(M)** Quantification of the 4-HNE-positive area fraction and TUNEL-positive apoptotic index in tumor sections. **(N)** Representative Ki-67 staining (top) and hematoxylin and eosin (H&E) staining (bottom) of tumor sections. Scale bar, 100 μm. **(O)** Quantification of Ki67-positive cells in tumor sections. Data are presented as mean ± SD. Statistical significance was determined by one-way ANOVA with Tukey’s multiple comparisons test. n = 5 mice per group for (C) to (G) and n = 3 mice per group for (H) to (O). \**P* < 0.05, \*\**P < 0.01,* \*\**\*P < 0.001,* \*\*\*\**P* < 0.0001.

Given that our *in vitro* data had already shown that this system induces ferroptosis-associated damage in tumor cells, we next examined whether comparable lipid peroxidation phenotypes and tumoricidal effects could also be observed *in vivo*. We performed 4-hydroxynonenal (4-HNE) staining to assess lipid peroxidation-associated oxidative damage in tumor tissues. 4-HNE levels were markedly increased in the G6 group compared to other groups, indicating elevated lipid peroxidation in tumor tissues (Fig. 6L). These findings, together with the increased intratumoral iron deposition, suggest that LMR-Immunobot treatment under HMF promotes ferroptosis-associated oxidative damage *in vivo*. Meanwhile, TUNEL-positive signals were significantly increased in G6 (Fig. 6L and M), indicating enhanced terminal cell death. In addition, compared with G1, G6 showed markedly reduced Ki-67 staining, revealing significant suppression of tumor cell proliferation, while H&E staining revealed more extensive necrotic regions and more severe tissue destruction (Fig. 6N and O). Collectively, these results demonstrate that the LMR-Immunobot + HMF strategy enhances ferroptosis-associated oxidative damage and tumor killing, thereby achieving the most potent antitumor efficacy *in vivo*.

To further evaluate the *in vivo* safety of this therapeutic strategy, we analyzed major organs and serum biochemical parameters. No obvious abnormal signs or significant body weight loss were observed during treatment (fig. S20). At the endpoint, H&E staining of major organs, including the hearts, livers, spleens, lungs, and kidneys, showed no apparent pathological damage, and serum biochemical analyses did not indicate apparent hepatic or renal dysfunction or widespread tissue toxicity (fig. S21). These findings suggest that immunobots exhibit favorable *in vivo* biocompatibility and systemic safety under the dosing and magnetic actuation conditions used in this study.

### Immunobots under hybrid magnetic actuation remodel TME and boost immune responses

Tumor-associated macrophages constitute a key myeloid component of the colorectal cancer immune microenvironment, and their polarization state is closely linked to tumor progression and therapeutic responsiveness (*45*, *46*). In general, M1-like macrophages are associated with proinflammatory and antitumor activities, whereas M2-like macrophages are more commonly implicated in immunosuppression, tissue remodeling, and tumor progression (*47*). Given that immunobots are built on a BMDM chassis, we next asked whether its intratumoral accumulation could also reshape the local macrophage niche. Flow cytometric analysis showed that the proportion of M1-like macrophages (CD80^+^CD86^+^ within F4/80^+^CD11b^+^ cells) was significantly increased from approximately 26% in the control group to 48% in tumors treated with the LMR-Immunobot + HMF group, whereas M2-like macrophages (CD206^+^ within F4/80^+^CD11b^+^ cells) were markedly reduced from ∼29% down to approximately 13% (Fig. 7A and fig. S22). Immunofluorescence staining further confirmed an increase in CD80^+^F4/80^+^ cells together with an accompanying decrease in CD206^+^F4/80^+^ cells (Fig. 7B and C and fig. S24A). Consistently, ELISA revealed elevated levels of TNF-α and IL-12, accompanied by reduced IL-10 levels (Fig. 7D). Together, these findings indicate that, upon tumor accumulation, immunobots drive TAM polarization toward a proinflammatory, antitumor state, thereby establishing a myeloid microenvironment more permissive for subsequent immune activation.

**Fig. 7.**
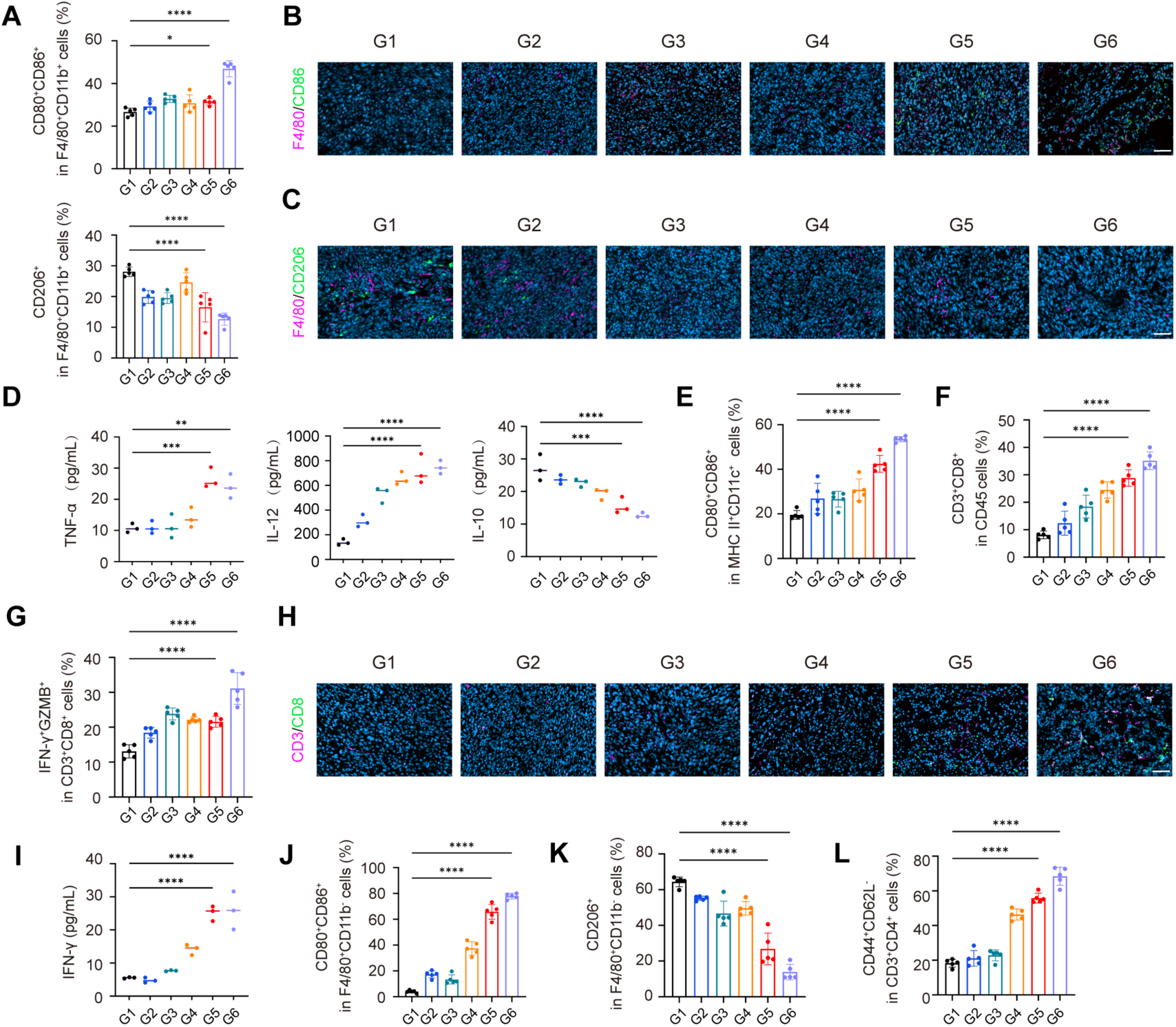
Analysis of tumor microenvironment and immune response. (**A**) Flow cytometric analysis of M1-like (CD80^+^CD86^+^) and M2-like (CD206^+^) macrophage populations in tumors from the indicated groups. (**B**) Representative immunofluorescence images of CD80 (green) and F4/80 (magenta) staining in tumor sections showing the increased accumulation of M1-macrophages in G6. Scale bar, 200 μm. (**C**) Representative immunofluorescence images of CD206 (green) and F4/80 (magenta) staining in tumor sections. Scale bar, 200 μm. (**D**) Serum levels of TNF-α, IL-12, and IL-10, measured by ELISA. (**E**) Flow cytometric analysis showing the proportion of mature dendritic cells in tumors of each group. (**F**) Flow cytometric analysis of CD3^+^CD8^+^ T cell population in tumors. (**G**) Flow cytometric quantification of IFN-γ^+^GZMB^+^ CD8^+^ T cells in tumors. (**H**) Representative immunofluorescence images of CD3 and CD8 staining in tumor sections further confirming the flow cytometric analysis. Scale bar, 200 μm. (**I**) Serum IFN-γ levels measured by ELISA. (**J** and **K**) Flow cytometric quantification of CD80⁺CD86⁺ M1-like (J) and CD206⁺ M2-like (K) macrophages within the splenic RPM-enriched population (CD45⁺F4/80⁺CD11b⁻ cells). (**L**) Flow cytometric analysis of CD44^+^CD62L⁻ cells gated on CD3^+^CD4^+^ splenocytes. Data are presented as mean ± SD. Flow cytometry data were obtained from 5 mice per group, and ELISA data were obtained from 3 mice per group. Statistical significance was determined by one-way ANOVA with Tukey’s multiple-comparison test.

Whether immunobot-induced myeloid remodeling translates into effective adaptive antitumor immunity depends in part on functional antigen presentation by mature dendritic cells (DCs). As professional antigen-presenting cells that bridge innate and adaptive immunity, mature DCs can process tumor-associated antigens and promote antitumor CD8⁺ T-cell responses through antigen presentation and cross-priming (*48*, *49*). On this basis, we next examined the effects of immunobots on DC maturation and CD8^+^ T cell activation. The LMR-Immunobot + HMF group showed a significant increase in mature DCs (CD80^+^CD86^+^ within MHC II^+^CD11c^+^ cells), rising from ∼20% to approximately 52% (Fig. 7E), together with enhanced infiltration of CD3^+^CD8^+^ T cells in tumors, which expanded from ∼8% in the control to approximately 34% (Fig. 7F and H and figs. S23 and S24B). More importantly, this group showed a marked increase in IFN-γ^+^GZMB^+^ cells within CD3^+^CD8^+^ T cells, jumping from ∼14% in the control to ∼30% (Fig. 7G), indicating enhanced cytotoxic effector function of intratumoral CD8^+^ T cells, along with a corresponding increase in serum IFN-γ levels (Fig. 7I). These results suggest that immunobot-induced myeloid reprogramming enhances antigen presentation and cytotoxic T cell responses, thereby promoting antitumor immunity within the tumor microenvironment.

To determine whether immunobot-induced immune remodeling extended beyond the tumor microenvironment, we next analyzed splenic immune composition as a peripheral readout of systemic immune alterations. Red pulp macrophages (RPMs), a major splenic tissue-resident macrophage population characterized by high F4/80 and low or absent CD11b expression, participate in blood-cell clearance, iron handling, and immune regulation (*50*). In the LMR-Immunobot + HMF group, the proportion of M1-like cells within the splenic RPM-enriched population (CD80⁺CD86⁺ within F4/80⁺CD11b⁻ cells) increased from approximately 8% to 78%, whereas the proportion of M2-like cells (CD206⁺ within F4/80⁺CD11b⁻ cells) decreased from approximately 65% to 15% (Fig. 7J and K and fig. S25). These results indicate a pronounced proinflammatory shift within the splenic resident macrophage compartment. We further examined the splenic CD4⁺ T-cell compartment. Within CD3⁺CD4⁺ T cells, CD44⁺CD62L⁻ cells represent an effector-memory phenotype, antigen-experienced population. This subset increased from approximately 20% to 70% following LMR-Immunobot + HMF treatment (Fig. 7L and fig. S26), indicating an expansion of peripheral antigen-experienced CD4⁺ T cells. Together with enhanced intratumoral DC maturation and cytotoxic CD8⁺ T-cell activation, these findings indicate that immunobot-mediated immune remodeling involves coordinated alterations in both local and peripheral immune compartments.

To provide transcriptomic support for the *in vivo* findings, we performed bulk RNA sequencing on tumor tissues from the G1, G5, and G6 groups. Principal component analysis (PCA) showed clear separation among the three groups, with G6 forming a distinct cluster, indicating that HMF further reshaped the tumor transcriptional landscape, and differential expression analysis identified extensive treatment-associated transcriptional changes (Fig. 8A and B). Functional enrichment analyses showed that G6 tumors exhibited coordinated activation of inflammatory and antigen-processing programs, including TNF/NF-κB signaling and antigen presentation, together with oxidative stress-, ferroptosis-, and iron transport-related programs (Fig. 8C to E). Consistent with these pathway-level changes, the expression profiles of inflammatory and myeloid-associated genes, redox stress-related genes, and iron-handling genes were markedly altered in G6 tumors (Fig. 8F to H). Together, these transcriptomic data corroborate the phenotypic and functional findings above, indicating that LMR-Immunobot + HMF treatment is associated with an immune-activated tumor microenvironment accompanied by enhanced ferroptosis-related oxidative stress and iron-metabolic remodeling in tumor tissues.

**Fig. 8.**
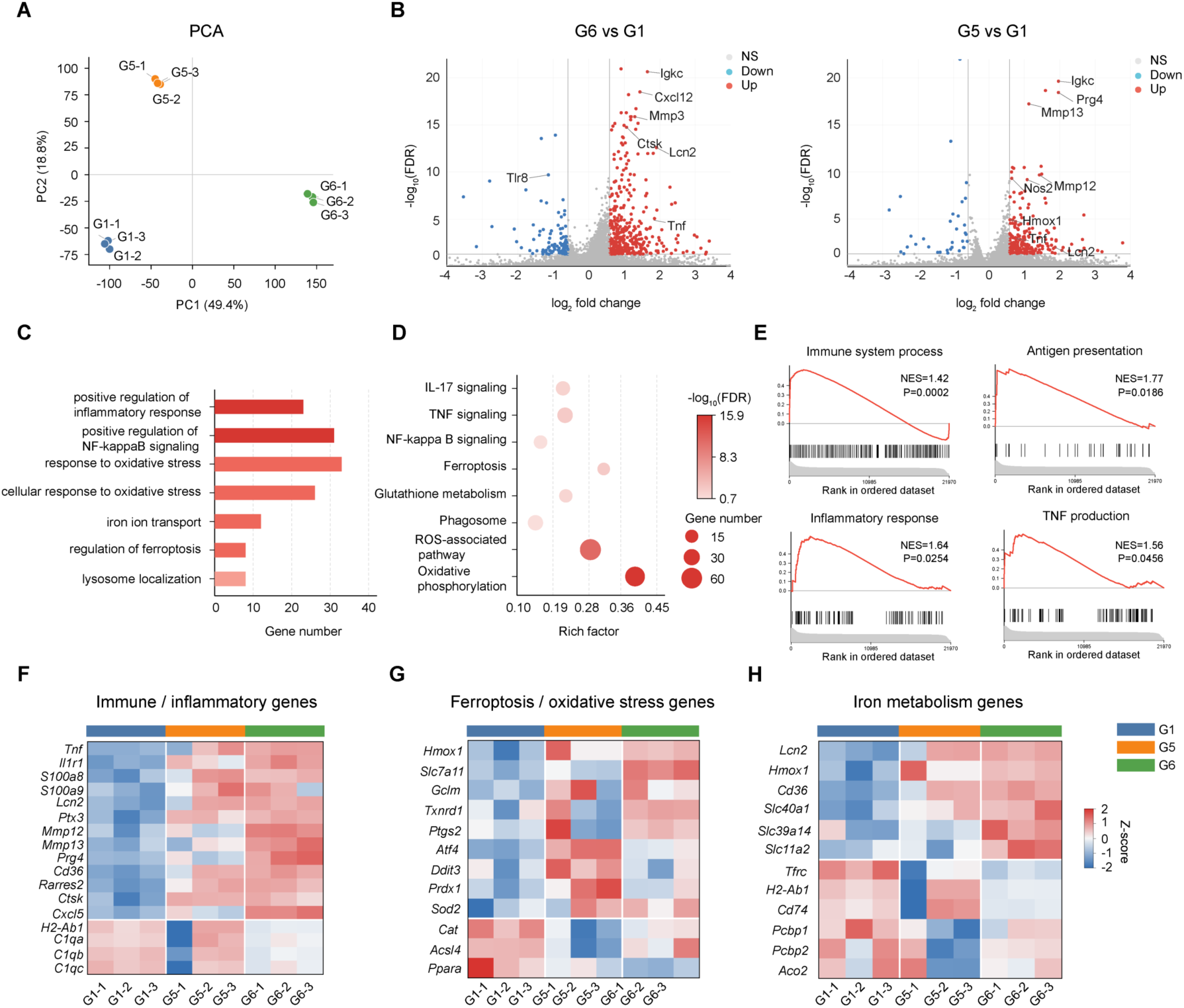
Transcriptomic profiling of tumor tissues after immunobot treatment. (**A**) PCA of bulk RNA-sequencing profiles from subcutaneous MC38 tumor tissues in the Saline (G1), LMR-Immunobot + GMF (G5), and LMR-Immunobot + HMF (G6) groups. Each dot represents one biologically independent tumor sample. (**B**) Volcano plots showing DEGs in G6 versus G1 and G5 versus G1 tumor tissues. Red and blue dots indicate significantly up-regulated and down-regulated genes, respectively, whereas gray dots indicate genes without significant differential expression. Representative altered genes are labeled. (**C**) GO enrichment analysis of DEGs in G6 versus G1 tumor tissues. (**D**) KEGG pathway enrichment analysis of DEGs in G6 versus G1 tumor tissues. Dot size indicates gene number, and color indicates −log_10_(FDR). (**E**) GSEA showing enrichment of immune system process, antigen presentation, inflammatory response, and TNF production signatures in G6 versus G1 tumor tissues. Normalized enrichment scores (NES) and *P* values are indicated in each plot. (**F** to **H**) Heatmaps showing row-scaled expression of representative immune/inflammatory genes (F), ferroptosis/oxidative stress-related genes (G), and iron metabolism-related genes (H) in tumor tissues from the G1, G5, and G6 groups. Red and blue indicate relatively higher and lower expression, respectively.

## Discussion

Solid tumor therapy remains limited by inefficient intratumoral transport of therapeutic agents and by an immunosuppressive microenvironment shaped by stromal, metabolic, and myeloid components (*11*, *51*). Macrophages are attractive cellular carriers because they can migrate toward and infiltrate tumor tissues. Their phenotypic plasticity, however, also limits their therapeutic application, because tumor-derived signals can drive macrophages toward tumor-supportive TAM-like states (*7*, *52*). In this study, we developed magnetically actuated macrophage-based immunobots to overcome these limitations. Internalized magnetic LMRs enabled external control of immunobot movement and improved tumor accumulation and tissue penetration. At the same time, LMR-derived iron together with LPS induced iron-metabolic remodeling in the carrier cells and activated ROS-NF-κB signaling, thereby promoting an M1-like antitumor program. These effects were associated with ferroptosis-related immunogenic tumor cell death and coordinated antitumor immune activation. Thus, immunobots provide a cellular therapeutic platform that integrates magnetically guided transport with local immunometabolic remodeling in solid tumors.

A key finding of this study is that the therapeutic transport capacity of immunobots depends on balancing magnetic payload with preserved cellular function. Previous macrophage microrobots have largely treated intracellular magnetic materials as enabling components for navigation, targeting, or drug delivery (*30*, *53–56*). More recent immunobot studies have further coupled magnetic microrollers with macrophage activation and imaging-guided control (*17*, *30*). However, how the amount of intracellular magnetic cargo affects the transport function of living carrier cells has remained insufficiently understood. Here, we show that distinct microroller loading states regulate not only magnetic propulsion but also the intrinsic migratory and barrier-penetrating capacities of macrophages. Increasing LMR loading enhanced magnetic responsiveness, whereas excessive loading impaired spontaneous macrophage motility and transbarrier penetration. Intermediate loading provided a more favorable balance by retaining effective magnetic control while preserving the cellular properties required for tumor infiltration. Notably, remagnetization after internalization further improved propulsion by restoring efficient torque transfer from intracellular hard-magnetic microrollers. These findings indicate that therapeutic transport in living microrobots should not rely on payload maximization alone, but rather on matching magnetic actuation with the intrinsic transport capacity of the carrier cell. Payload heterogeneity generated during macrophage-mediated assembly remains an engineering challenge, as immunobots with different loading states may exhibit distinct motility profiles that affect population-level synchronization and control. By tuning the microroller-to-cell ratio, we enriched single- and dual-loaded immunobots and preserved a favorable balance between magnetic actuation and intrinsic macrophage motility. Future work incorporating microfluidic or chip-assisted assembly could further reduce payload heterogeneity and improve population-level actuation.

Immunobot-induced tumor ferroptosis further suggests a functional relationship between macrophage iron handling and tumor-cell metabolic vulnerability. Inflammatory M1 macrophages preferentially adopt an iron-withholding phenotype associated with increased ferritin heavy-chain expression and reduced transferrin receptor expression relative to M2 macrophages (*19*). Consistent with this phenotype, BMDMs that internalized LMRs accumulated intracellular iron without undergoing ferroptosis and displayed increased FTH1 and reduced TFR1 expression. These findings suggest that carrier macrophages can buffer LMR-derived iron while maintaining inflammatory activation and viability. Tumor cells exposed to immunobots showed a different response, with decreased FTH1, increased TFR1, and ferroptotic death. Because TFR1 is associated with ferroptotic cells and reduced ferritin buffering is expected to increase susceptibility to labile iron accumulation, these findings support a cell-context-dependent response to the iron-containing platform rather than nonspecific iron toxicity (*57*). However, the route by which LMR-derived iron becomes available to tumor cells remains unresolved. Based on our transcriptomic and biochemical data, we speculate that this process may involve cooperation between inflammatory stress and FPN-independent iron transfer. Immunobots may weaken tumor antioxidant defenses through TNF-α, IL-12, Nos2, and Il1b signaling, while LMR-derived iron may be released into the local microenvironment through ferritin- or iron complex-containing extracellular vesicles, secretory lysosome/autophagy-associated noncanonical ferritin secretion, or iron-rich extracellular particles and cell fragments. Under conditions of increased TFR1 expression and impaired FTH1 buffering in tumor cells, these iron inputs may be more readily converted into labile iron accumulation and lipid peroxidation. This model will require further validation by high-resolution iron tracing, EV secretion blockade, perturbation of ferritin secretion pathways, and inhibition of iron uptake routes.

Ferroptosis in this system may also function as an immune-amplifying event. Immunobot treatment increased surface CRT exposure and extracellular ATP and HMGB1 release from tumor cells, whereas these ICD-associated signals were reduced by Fer-1 treatment, establishing a functional connection between ferroptotic tumor-cell death and the induction of an immunogenic phenotype. Surface-exposed CRT serves as a pro-phagocytic signal that promotes the recognition and engulfment of dying or malignant cells by macrophages through the CRT-LRP/CD91 axis (*43*). Consistent with this pro-phagocytic mechanism, immunobot treatment enhanced the engulfment of tumor cells by macrophages, whereas Fer-1 treatment of tumor cells markedly reduced their subsequent uptake by immunobots. These findings suggest that ferroptosis-associated immunogenic signals contribute to immunobot-mediated phagocytic clearance of tumor cells. Separately, immunobot treatment generated a proinflammatory tumor microenvironment and was accompanied by a shift of resident TAMs toward an antitumor phenotype. Whether these changes in the myeloid compartment are translated into effective adaptive antitumor immunity depends in part on mature dendritic cells, which process tumor-associated antigens and promote CD8⁺ T-cell responses through antigen presentation and cross-priming (*48*). In agreement with this framework, immunobot treatment increased intratumoral DC maturation and cytotoxic CD8⁺ T-cell activation. Activated CD8⁺ T cells may in turn further sensitize tumor cells to ferroptosis, as IFN-γ has been shown to suppress SLC7A11 and SLC3A2 expression, restrict cystine uptake, and promote lipid peroxidation in tumor cells (*24*). Nevertheless, the immune consequences of ferroptosis are context dependent, because ferroptotic cancer cells can impair dendritic cell maturation and antigen cross-presentation under certain conditions (*27*). Together, these findings are consistent with a ferroptosis-associated immune amplification process involving immunogenic tumor-cell death, myeloid remodeling, antigen presentation, and cytotoxic T-cell activity.

The broader significance of this work lies in integrating magnetic navigation with the exploitation of metabolic vulnerability and antitumor immune activation, three therapeutic principles that are often pursued separately. Ferroptosis-inducing nanomedicines offer a promising means of exploiting tumor redox vulnerability, but their efficacy and safety depend on achieving sufficient delivery within solid tumors while limiting iron-dependent oxidative damage in normal tissues (*11*). Against this background, immunobots provide a cell-based strategy for improving tumor delivery. Magnetic actuation enhanced tumor accumulation and penetration, while iron buffering in carrier macrophages preserved cell viability and supported an inflammatory M1-like phenotype. Within the treated tumor microenvironment, enhanced tumor-cell ferroptotic susceptibility was accompanied by DC maturation and CD8⁺ T-cell activation, indicating coordinated metabolic and immune remodeling. Future studies should determine whether the local metabolic and immune remodeling induced by immunobots can generate durable systemic antitumor immunity and remain effective across tumors with distinct stromal and immune architectures. Overall, this study supports a therapeutic framework in which living microrobots combine directed tumor delivery with localized metabolic intervention and immune activation.

## Materials and Methods

### Janus FePt microroller fabrication

The Si (100) substrate was cleaned with deionized water and ethanol by ultrasonication followed by oxygen plasma treatment. A commercial silicon dioxide microsphere solution (5 μm, Sigma Aldrich) was then drop-cast onto the treated wafers and formed a uniform monolayer distribution. To deposit FePt films on top of SiO₂ particles, samples were loaded into an Explorer 18 DC magnetron sputtering system (Denton Vacuum, LLC) with a background vacuum maintained below 3 × 10^-6^ Pa. Both Fe and Pt targets had a purity of 99.99%, and the argon working pressure was 0.67 Pa. The composition of the samples was controlled by adjusting the sputtering power of the two targets. The film thickness was regulated by modifying the total sputtering time. After sputtering, the samples were annealed in a tube furnace at 650 °C for 1 h under vacuum conditions to achieve a stable L1_0_ phase characterized by high remanence and coercivity.

### Characterization of FePt microrollers

An impulse magnetizer (DPM1, EUSCI) was used to magnetize the FePt film with an out-of-plane direction. The FePt-coated silica particles were collected in a sterile PBS solution via sonication for subsequent experiments. The surface morphology of FePt films was examined by SEM (MIRA4, TESCAN). EDS (BRUKER XFlash 630m) was used to assess the uniformity of the films and the spatial distribution of Fe and Pt in the coating. The exact ratios of Fe and Pt contained in the samples were determined by Inductively Coupled Plasma Optical Emission Spectroscopy (Agilent Technologies 5110). XRD measurements were performed with Empyrean 3.0, Malvern PANalytical, using a sealed tube Cu-anode with a 0.05° scanning step size. The magnetic properties of microrollers with different FePt atomic ratios were measured with a vibrating sample magnetometer (Lake Shore 7404) in a ±2 T magnetic field.

### Fabrication of LPS-modified FePt microrollers

(3-aminopropyl) triethoxysilane (APTES, 98%, Macklin) was introduced to graft primary amine groups onto the silica hemisphere. The microrollers were dispersed in ethanol (1 mL) and mixed with 50 µL of APTES under vortex agitation for 3 h at 37 °C. The microrollers were then collected by centrifugation and thoroughly rinsed with ethanol and dimethyl sulfoxide (DMSO, Biosharp). In the next step, surface amine groups were reacted with N-hydroxysuccinimide (NHS)-biotin via NHS-amine coupling. Biotin NHS (5 mg/mL, MCE) was added to the particle suspension and gently vortexed for 3 h. After completion of the reaction, excess reagents were removed by repeated washing with Dulbecco’s phosphate-buffered saline (DPBS, Macklin). Subsequently, streptavidin-Cy3 (APExBIO) was conjugated to the biotinylated particles through the high-affinity biotin–streptavidin interaction. The particles were incubated with a streptavidin solution (50 µg/mL in DPBS) for 1 h, followed by extensive washing with DPBS. Finally, biotinylated LPS derived from Escherichia coli O55: B5 (InvivoGen) was immobilized on the particle surface using the same biotin–streptavidin binding strategy. The particle suspension was incubated with LPS (100, 200, 500 µg/mL in PBS) for 1 h and subsequently washed with DPBS to eliminate unbound molecules. The functionalized magnetic microrollers were resuspended in DPBS and sterilized under ultraviolet light before further use.

### Zeta potential measurements

The surface charge alterations of the 5 µm bare SiO₂ microspheres, FePt-coated Janus microrollers (MRs), and LPS-functionalized microrollers (LMRs) were evaluated via zeta potential analysis. Prior to measurement, the respective microparticle samples were thoroughly washed and uniformly dispersed in PBS (pH 7.4) to maintain physiological conditions. Measurements were conducted at room temperature using a dynamic light scattering instrument (Zetasizer Pro, Malvern Panalytical). All tests were performed in triplicate, and data were recorded to calculate the average values and standard deviations.

### Cell culture and maintenance

All cell types were cultured in proliferation medium (DMEM supplemented with 10% fetal bovine serum and 1% penicillin-streptomycin, all from Gibco) and maintained at 37 °C in a humidified incubator with 5% CO₂. The medium was refreshed every two days, and cells were subcultured or harvested for subsequent experiments when they reached 80 to 90% confluence. Among them, the mouse colon carcinoma MC38 cell line and human umbilical vein endothelial cells (HUVECs) were kindly provided by the laboratory of Professor Mei Song at Sun Yat-sen University.

### Isolation and culture of mouse BMDMs

For the collection of BMDMs, mononuclear cells were isolated from the femurs of 6-week-old male mice and cultured in proliferation medium containing 50 ng/mL macrophage colony-stimulating factor (M-CSF, MCE, HY-P7085). The medium was replaced every two days, and non-adherent cells were removed to enrich the macrophage population. After 1week, mature macrophages were collected and subjected to a 24 h induction period for M1 polarization (using 100 ng/mL LPS, Sigma, L2880), M2 polarization (using 20 ng/mL IL-4, PeproTech, 214-14-20UG), or microroller stimulation. The stimulated BMDMs were then used for subsequent experiments.

### CCK-8 assay

BMDMs were seeded in 96-well plates at 3 × 10⁵ cells per well and allowed to adhere overnight. Cells were then incubated with LMRs at 0.05 mg mL⁻¹ for 24, 48, or 96 h. Cell viability was measured using a CCK-8 assay kit (Yeasen) according to the manufacturer’s instructions. Briefly, 10 μL of CCK-8 reagent was added to each well containing 100 μL of medium and incubated at 37 °C with 5% CO₂. Absorbance was measured at 450 nm using a microplate reader.

### Magnetic steering of immunobots

The magnetic microrollers were actuated by a custom-built five-coil electromagnetic system (four xy coils and one z coil), integrated with an inverted bright-field microscope (CKX53, 389 Olympus). The step-out frequency of the microrollers was determined by gradually increasing the frequency of the rotating magnetic field and finding the maximum frequency at which the microrollers could no longer follow. Similar experiments were performed in a closed microchannel containing a monolayer of endothelial cells, simulating a blood-vessel flow environment. The mean speeds and trajectories of microrollers were analyzed using a customized Python code that performs object detection and tracking.

### Magnetic capture of immunobots under flow

Microfluidic channels were fabricated by PDMS casting. Briefly, PDMS prepolymer and curing agent were mixed at a 10:1 ratio, poured into a CNC-machined high-temperature-resistant resin mold, and cured at 70 °C for 1 h. The cured PDMS chip was gently removed, plasma-treated, and bonded to a glass slide.

To assess magnetic capture under flow, immunobots were suspended in serum and perfused through a vessel-mimetic cylindrical microchannel with an inner diameter of 300 μm. Linear flow velocities were set to 1.8, 3.6, and 7.2 mm s⁻¹ using a syringe pump (JSP-02-1C, Mifluidic), and magnetic capture was tested under field strengths of 10, 50, and 250 mT. A permanent magnet was positioned adjacent to the channel to generate a transverse magnetic field gradient and attract immunobots toward the magnet-facing channel wall. Fluorescence videos were acquired at the initial, intermediate, and final time points using a high-resolution fluorescence stereo microscope (M205 FA, Leica). Immunobot accumulation was quantified as the fluorescence-positive area within the magnet-facing region of interest. Final retention was normalized to the initial signal in the same channel as (At_final_ - At_0_) to correct for loading heterogeneity. Estimated wall shear stress was calculated as equation (1):

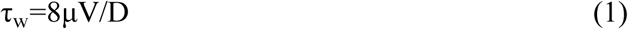

where μ is the serum dynamic viscosity, V is the mean linear velocity, and D is the channel diameter.

### Viability after continuous magnetic actuation

Immunobots were first subjected to continuous magnetic actuation under a 10 mT rotating magnetic field at 30 Hz for 15 min. After magnetic actuation, the immunobots were incubated for different durations, followed by cell viability assessment using a CCK-8 assay.

### Transwell assay

The *in vitro* penetration capability of immunobots was evaluated using 8 µm pore-size transwell inserts (Corning Inc.). Approximately 10,000 CFSE-labeled BMDMs or immunobots were seeded in the upper chamber with DMEM in the lower chamber. Before magnetic field application, multiple random fluorescence micrographs were acquired from the upper chamber to determine the initial distribution of single-, dual-, and multi-payload immunobots. The fraction of each payload category was calculated from these images and used to estimate the input number of each group according to following equation (2):

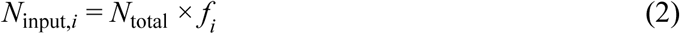

where *N*_total_ is the total number of immunobots seeded in the upper chamber and *f*_i_ is the initial fraction of payload category i. After transmigration, the number of single-, dual-, and multi-payload immunobots in the lower chamber was quantified by fluorescence microscopy. The payload-specific penetration rate was calculated as equation (3):

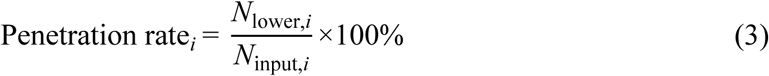

A 250 mT gradient magnetic field or a hybrid magnetic field consisting of this gradient magnetic field and a 10 mT, 20 Hz rotating magnetic field was applied beneath the transwell plate for 1 h. The migrated cells in the lower chamber were observed by optical microscopy and quantified by fluorescence intensity using a microplate reader.

For tumor cell-related experiments, M0, M1 macrophages, or immunobots were seeded in the upper chamber of 0.4 µm transwell inserts (Corning), while MC38 cells were seeded in the lower chamber. Cells were pretreated with DFO (50 µM; MCE, HY-B0988), erastin (10μM; Selleck, S7242), or Fer-1 (5μM; Selleck, S7243) for 1 h before coculture, followed by 24 h of coculture before subsequent analyses.

### LMR degradation kinetics and magnetic stability evaluation

Fe release from LMR was assessed in phagolysosomal simulant fluid (PSF, pH 4.5) at 37 °C. For each time point (12 h, 1, 3, 5, and 7 days), an independently prepared particle aliquot was incubated in PSF under gentle agitation, followed by centrifugation to separate the supernatant and residual microrollers. The collected supernatant was acidified and analyzed by ICP-MS (Inductively Coupled Plasma Mass Spectrometry) to determine the released Fe content. The corresponding particle pellet was subsequently fully digested in nitric acid and also analyzed by ICP-MS (5110 ICP-OES, Agilent Technologies) to quantify the remaining Fe. The Fe release percentage was calculated as the following equation (4):

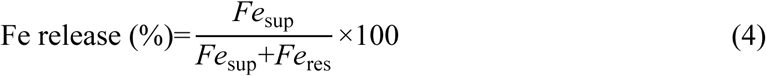

This calculation normalized the result to the total Fe recovered from each individual tube and minimizing errors arising from small differences in initial particle loading. Blank PSF controls were included for background correction. The residual MRs were collected and subjected to VSM analysis to confirm the structural and magnetic retention of the L1_0_-FePt core.

### Flow cytometry

For cellular flow cytometry, cells were collected after the indicated treatments, washed with cold PBS, and stained for flow cytometry analysis. For macrophage polarization, cells were incubated with APC-conjugated anti-CD86 or anti-CD206 antibodies at 4 °C for 30 min in the dark. Intracellular ROS levels were detected using a DCFH-DA probe (10 μM; Yeasen, Cat. No. 50101ES01), and cell death was assessed by Annexin V/PI staining (Beyotime, Cat. No. C1062S) according to the manufacturers’ instructions.

For tissue flow cytometry, mice were euthanized at the experimental endpoint to analyze tumor-infiltrating and splenic immune cells. Tumors were minced and digested with 1 mg mL⁻¹ collagenase IV (Gibco), 0.1 mg mL⁻¹ DNase I (Sigma), and 0.1 mg mL⁻¹ hyaluronidase (Sigma) at 37 °C for 45 min, followed by filtration through a 70 μm cell strainer. Spleens were mechanically dissociated through a 70 μm cell strainer, and red blood cells were lysed with 1× RBC lysis buffer (BioLegend). Single-cell suspensions were blocked with anti-mouse CD16/CD32 antibody for 10 min on ice, stained with Zombie Fixable Viability Dye to exclude dead cells, and surface-stained with fluorochrome-conjugated antibodies against CD45, CD11b, F4/80, CD80, CD86, CD11c, MHC II, CD3, CD4, CD8a, CD44, and CD62L (table S2) at 4 °C for 30 min. For intracellular staining, tumor single-cell suspensions were re-stimulated for 4 h at 37 °C in complete RPMI-1640 medium containing Cell Activation Cocktail without Brefeldin A (BioLegend, Cat. No. 423301), Brefeldin A Solution (BioLegend, Cat. No. 420601), and Monensin Solution (BioLegend, Cat. No. 420701), followed by fixation/permeabilization using a BD Cytofix/Cytoperm Kit and staining with antibodies against CD206, IFN-γ, and Granzyme B. Samples were acquired on a BD FACSymphony A1 flow cytometer and analyzed using FlowJo software v10.8.1 (BD Biosciences).

### ELISA

Serum and cell culture supernatant levels of inflammatory cytokines and HMGB1 were measured using mouse ELISA kits according to the manufacturers’ instructions. The analytes included TNF-α (GBA, CME0004-G048), IL-6 (GBA, CME0006-G048), IL-10 (GBA, CME0016-G048), IL-12/23(p40) (GBA, CME0024-G048), IFN-γ (GBA, CME0003-G048), and HMGB1 (Elabscience, E-EL-M0676). Absorbance was measured at 450 nm, and concentrations were calculated using standard curves.

### Cellular fluorescence staining

For the detection of intracellular Fe²⁺, cells from different treatment groups, including Immunobot, Immunobot + FAC (50 µg/ml; MCE, HY-B1645), and Immunobot + DFO (50 µM; MCE, HY-B0988), were washed with phosphate-buffered saline (PBS) and subsequently incubated with 1 µmol/L Ferro Orange probe (Dojindo, F374-10) in a serum-free medium. After a 30-min incubation at 37 °C in the dark, the cells were directly subjected to imaging.

Cell viability was evaluated using a Live/Dead cell imaging kit (Thermo Fisher Scientific, L3224). Following the treatments, the cells were washed with PBS and incubated with the staining working solution prepared strictly according to the manufacturer’s instructions. After a 30-min incubation at room temperature in the dark, the cells were washed twice with PBS and immediately subjected to fluorescence imaging.

Intracellular lipid peroxidation (LPO) levels were measured using the BODIPY™ 581/591 C11 probe (MCE, HY-DY1022) according to the manufacturer’s protocol. The cells were incubated with the freshly prepared dye working solution, which was supplemented with Hoechst for nuclear counterstaining, for 30 min at 37 °C in the dark. Prior to imaging, the cells were washed three times with PBS to remove excess dye.

For CRT immunofluorescence, the treated cells were fixed with 4% paraformaldehyde (PFA) for 15 min at room temperature and then blocked with 5% bovine serum albumin (BSA) for 1 h at room temperature. Subsequently, the cells were incubated with a specific primary anti-CRT antibody overnight at 4 °C. On the following day, the cells were washed with PBST and incubated with a FITC-conjugated secondary antibody for 1 h at room temperature in the dark. Finally, the nuclei were counterstained with DAPI. All fluorescence images were acquired using a laser scanning confocal microscope (LSM 980, Carl Zeiss), and quantitative analysis of the fluorescence intensity was performed using ImageJ software.

### Western blotting analysis

Proteins extracted in RIPA buffer with protease/phosphatase inhibitors were quantified by BCA assay. Equal protein aliquots (20 μg) were resolved by SDS-PAGE and transferred to PVDF membranes. Membranes were blocked (5% milk or 3% BSA, 1 h), probed with primary antibodies (overnight, 4 °C) and secondary antibodies (1 h, room temperature), and visualized using ECL on a Bio-Rad system. Band intensities were quantified via ImageJ.

### MDA assay

Lipid peroxidation was evaluated by measuring malondialdehyde (MDA) levels using a Lipid Peroxidation MDA Assay Kit (Beyotime, Cat. No. S0131S). After the indicated treatments, cells or tumor tissues were collected, lysed or homogenized, and processed according to the manufacturer’s instructions. The absorbance was measured using a microplate reader, and MDA levels were calculated according to the standard curve.

### GSH/GSSG assay

Intracellular glutathione levels were detected using a GSH and GSSG Assay Kit (Beyotime, Cat. No. S0053). Briefly, cells were collected after the indicated treatments, lysed or homogenized, and subjected to GSH and GSSG detection according to the manufacturer’s protocol. The GSH/GSSG ratio was calculated to evaluate cellular redox status.

### ATP assay

Cellular ATP levels were measured using an Enhanced ATP Assay Kit (Beyotime, Cat. No. S0027). After treatment, cells were lysed, and ATP levels were determined according to the manufacturer’s instructions. Luminescence signals were detected using a microplate reader, and ATP concentrations were calculated based on the ATP standard curve.

### Live-cell imaging assay

MC38 cells were labeled with CFSE, and LMRs were conjugated with Cy3. To generate immunobots, BMDMs were incubated with Cy3-labeled LMRs for 24 h. The resulting immunobots were then cocultured with CFSE-labeled MC38 cells for 24 h. Live-cell imaging was performed using an Incucyte SX5 live-cell analysis system, and bright-field, green fluorescence, and red fluorescence images were acquired at the indicated time points.

### Animals and tumor model

C57BL/6J mice (male, 6 weeks old, 18 to 20 g) were purchased from the Laboratory Animal Facility of HKUST(GZ). All animal experiments were approved by the HKUST(GZ) Institutional Animal Care and Use Committee (IACUC, approval number: HKUST(GZ)-IACUC-2025-A0139). A total of 2 × 10^6^ MC38 cells were injected into the right hind limb of mice to establish a subcutaneous colorectal cancer tumor model. Tumor volume was measured by a vernier caliper and calculated according to the formula: V = L × W²/2, where L and W are the length and width of the tumor, respectively.

### *In vivo* magnetic guidance tumor targeting and biosafety assay

Mice were anesthetized with isoflurane during IVIS imaging. Mice in G4 and G5 received gradient magnetic guidance twice on day 0 and twice on day 1, with each session lasting 30 min. Mice in G6 received two rounds of hybrid magnetic actuation on day 0 and day 1; each round consisted of 30 min of gradient magnetic guidance followed by 30 min of rotating magnetic field exposure (15 Hz). IVIS imaging was performed after the daily magnetic treatments on days 0 and 1, and again on days 2, 4, and 6 to monitor tumor targeting and fluorescence clearance *in vivo*. Body weight was recorded on days 0, 1, 2, 4, 6, and 10. On day 10, mice were euthanized, and the hearts, livers, spleens, lungs, kidneys, and tumors were harvested for *ex vivo* IVIS imaging. Major organs were further collected for histological analysis, and serum was collected for biochemical assays to evaluate systemic biosafety. To evaluate the difference in immunobot accumulation within tumor tissue, ICP-MS was used to quantify the Pt concentration.

### Histology and image analysis

Tumor tissues were harvested, fixed in 4% paraformaldehyde, dehydrated in graded sucrose, embedded in OCT, and cryosectioned (10 μm). Tissue sections were stained with Prussian blue (Servicebio, G1029-3), hematoxylin and eosin (H&E; Servicebio, G1005), or TUNEL reagents (Beyotime, C1089) according to the manufacturers’ protocols. For immunofluorescence staining, cryosections were subjected to antigen retrieval and treated with 3% H_2_O_2_ to quench endogenous peroxidase activity, followed by blocking with 3% BSA (Servicebio, GC305006). Sections were incubated with primary antibodies against CD31, 4-HNE, Ki-67, CD86, CD206, F4/80, CD3, or CD8 overnight at 4 °C. After washing, sections were incubated with HRP-conjugated secondary antibody (Servicebio, GB23303) for 1 h at room temperature and developed using TSA reagent (Servicebio, G1226) for 10 min. When multiplex staining was required, sequential rounds of staining were performed on the same section with intermediate antibody stripping between cycles. Sections were counterstained with DAPI (Servicebio, G1012), mounted, and imaged using an automated slide scanning system. Fluorescence signals were quantified using ImageJ software on one section from each tumor, with three biologically independent tumors analyzed per group.

### RNA-seq analysis

Total RNA from cultured cells or tumor tissues was subjected to rRNA depletion and library construction for mRNA sequencing. Raw reads were filtered with fastp, aligned to the mouse reference genome (Ensembl release 115) using HISAT2, and quantified with StringTie and RSEM. Differential expression analysis was performed using DESeq2. Genes with FDR < 0.05 and |log₂ fold change| > 1 for cultured cells, or FDR < 0.05 and |log₂ fold change| > 0.585 for tumor tissues, were defined as differentially expressed genes. Heatmap, GO, KEGG, and GSEA were used for downstream pathway analyses.

### Statistical analysis

All experiments were repeated at least three times independently for reproducibility. Data are presented as mean ± SD. GraphPad Prism 10 software was used for the statistical analyses. Student’s t-test (individual group comparisons), one-way ANOVA (comparison of three or more groups), and two-way ANOVA (multiple experimental groups with several time points) were used to analyze statistical differences. *P* < 0.05 was considered statistically significant.

## Author contributions

Conceptualization: I.C.Y. Methodology: X.Z., Y.Z., and C.W. Investigation and validation: X.Z., Y.Z., C.W., M.F., Q.D., and K.W. Formal analysis: X.Z. and Y.Z. Data curation: X.Z. and Y.Z. Supervision: S.C. and I.C.Y. Funding acquisition: I.C.Y. Writing-original draft: X.Z., Y.Z., and I.C.Y. Writing-review and editing: all authors.

## Data and materials availability

All data needed to evaluate the conclusions in the paper are present in the paper and/or the Supplementary Materials. Additional data are available from the corresponding author upon reasonable request.

