## Supplementary materials for "Magnetically guided macrophage immunobots coordinate iron-metabolic cascades and immunogenic ferroptosis for tumor immunotherapy"

**This PDF file includes:**

Figs. S1 to S26
Tables S1 and S2
Legends for movies S1 to S7

**Other Supplementary Material for this manuscript includes the following:**

Movies S1 to S7

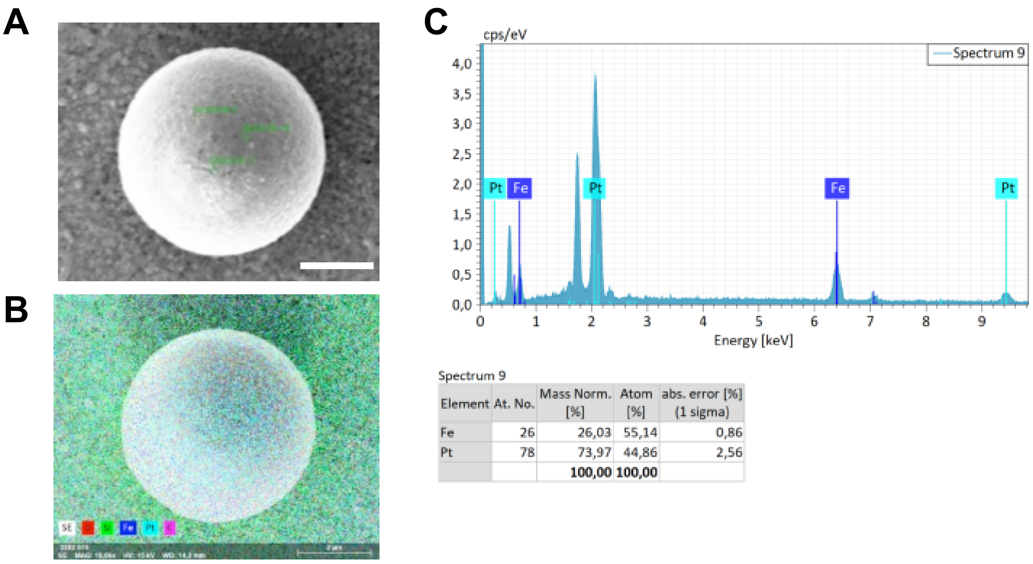


Spectrum 9

| Element | At. No. | Mass Norm. [%] | Atom [%] | abs. error [%] (1 sigma) |
|---|---|---|---|---|
| Fe | 26 | 26,03 | 55,14 | 0,86 |
| Pt | 78 | 73,97 | 44,86 | 2,56 |
| | | 100,00 | 100,00 | |

**Fig. S1.** Morphological and elemental characterization of FePt-coated microrollers. (A) Representative SEM image with EDS point scanning showing the positions, (B) element mapping of a single microroller showing the Fe and Pt localization on the surface. (C) EDS spectra showing the Fe/Pt mass percentage and atomic ratio of each micro-zone. Scale bar, 2 µm.

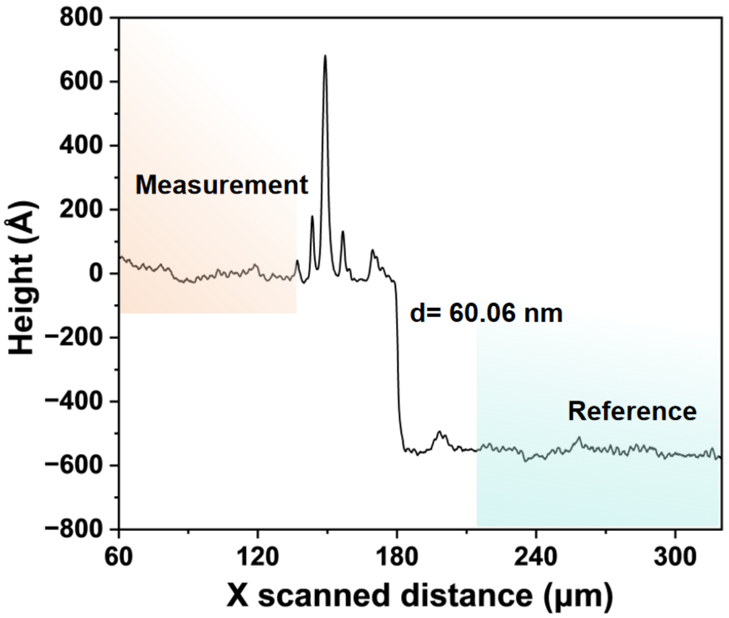


**Fig. S2.** Step profilometry measurement showing that the Fe200-Pt140 film thickness is 60.06 nm.

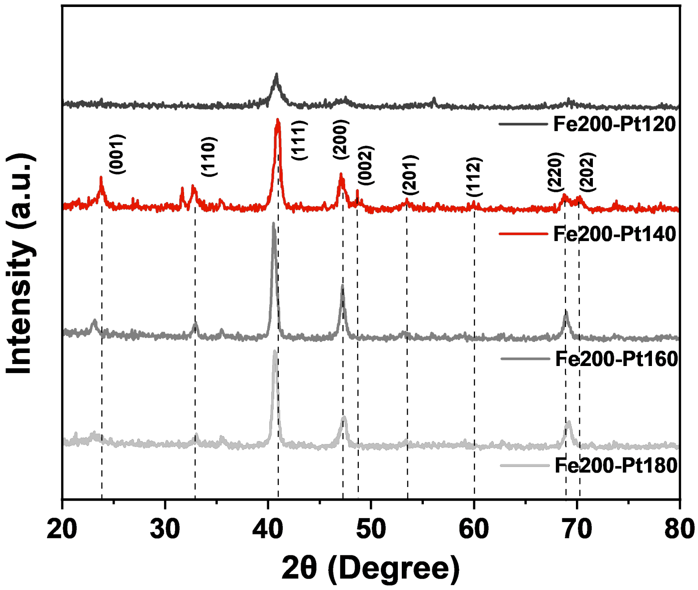


**Fig. S3.** XRD patterns of FePt films prepared under different Fe/Pt power ratios after annealing. Representative L10-FePt reflections are indicated in the Fe200-Pt140 group.

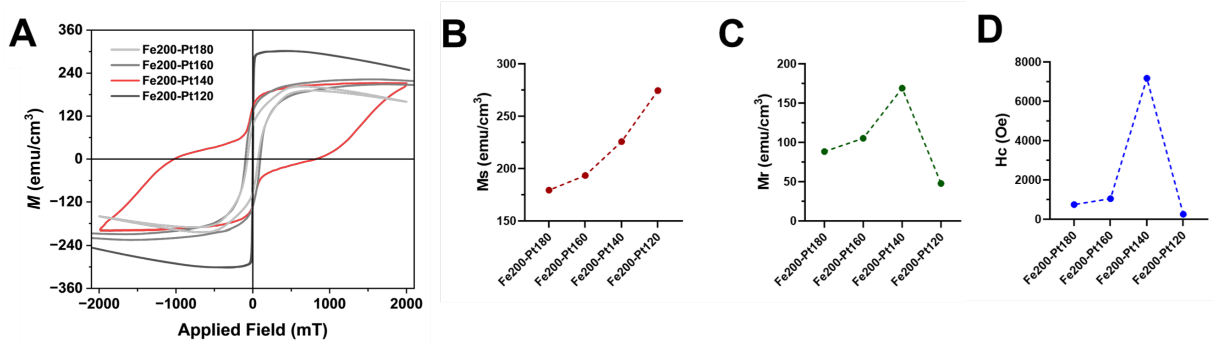


**Fig. S4.** Characterization of magnetic properties of the coated FePt using VSM. Hysteresis loops for different sputtering power groups. (A) Merged hysteresis loops. Changes in (B) Ms (saturation magnetization), (C) Mr (remanence) and (D) Hc (coercivity) among different groups after altering the FePt power ratio.

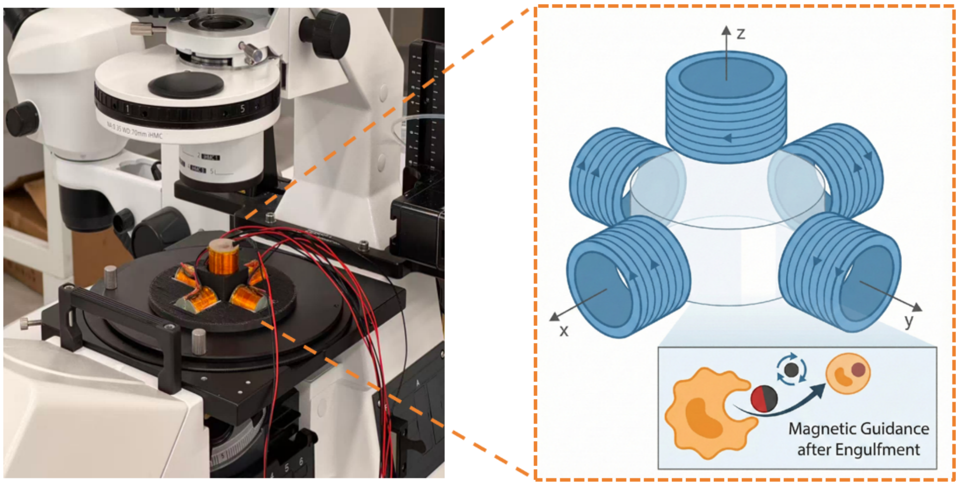


**Fig. S5.** Electromagnetic actuator setup used for *in vitro* actuation and magnetic navigation studies. Custom-built five-coil electromagnetic actuator system integrated with an inverted optical microscope for microroller and immunobot actuation. Inset showing a schematic illustration of each coil position and the sample space.

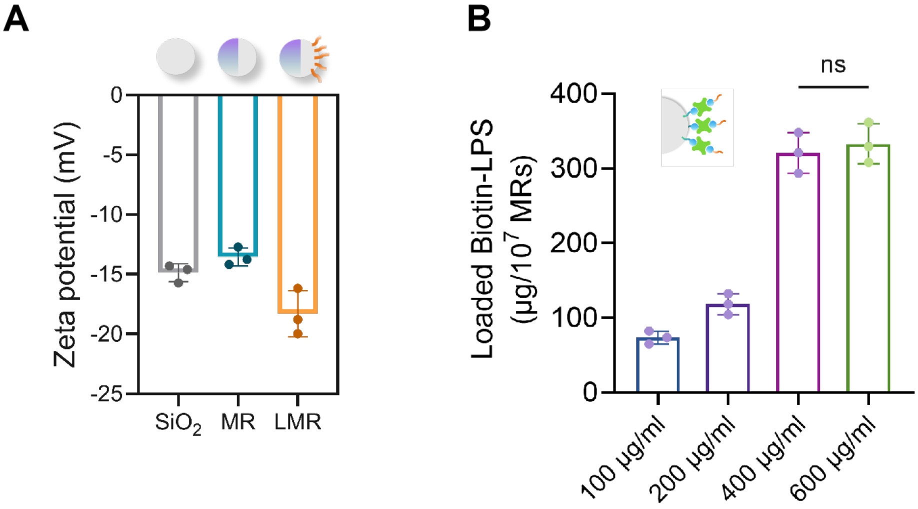


**Fig. S6. Physicochemical validation of LPS-engineered Janus FePt microrollers.** (A) Zeta potential alterations of the bare 5 μm $SiO_2$ microspheres, FePt-sputtered Janus microrollers (MR), and LPS-engineered microrollers (LMR) dispersed in PBS (pH 7.4). The distinct negative shift in the LMR group indicates the successful introduction of negatively charged LPS molecules onto the microroller surface. (B) Loading capacity of biotin-LPS on Janus microrollers dependent on initial LPS input. Data are presented as mean ± SD (n = 3).

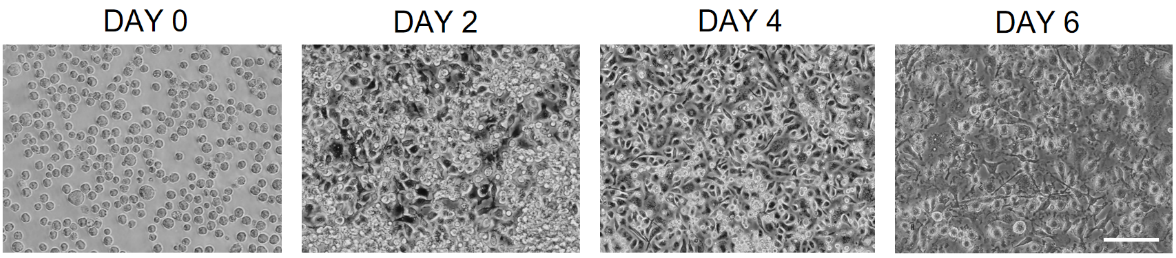


**Fig. S7. Differentiation of bone marrow progenitor cells into macrophages.** Morphological evolution of the cells at day 0, 2, 4, and 6 during the *in vitro* differentiation process. Scale bar, 50 μm.

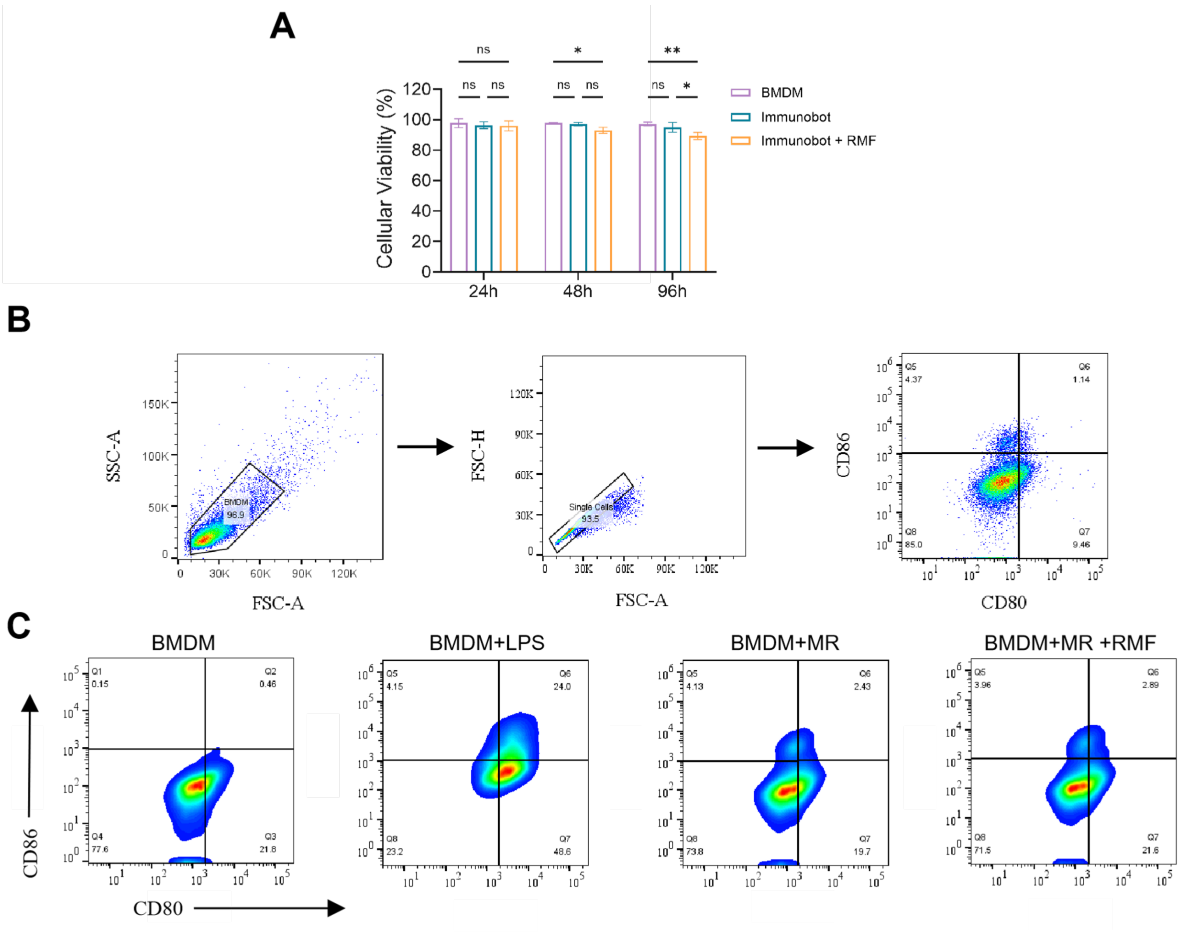


**Fig. S8. Effect of external magnetic rotation and steering on cell viability and phenotype.** (A) Viability results after rotating and steering at the step-out frequency (30 Hz) for 30 min. (B to C) FCM gating strategy and results indicating continuous rotation under RMF (30 Hz, 30 min) has a negligible effect on BMDM phenotype.

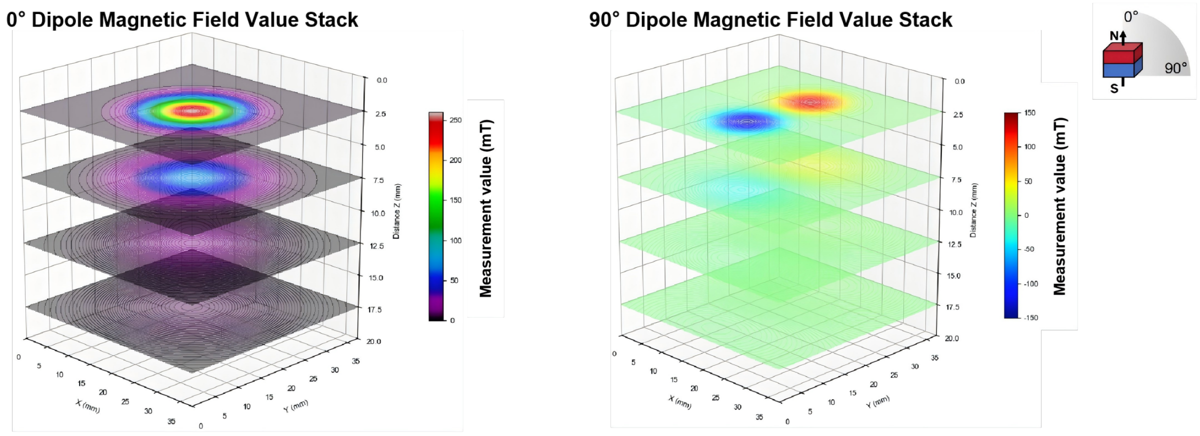


**Fig. S9. Effective magnetic field strength of the permanent magnets as a function of distance.** This magnet was used for the immunobots' flow-resistant retention and transwell penetration experiments. The figure shows the magnet at two angles over distance ($Z_{max}$ = 17.5 mm).

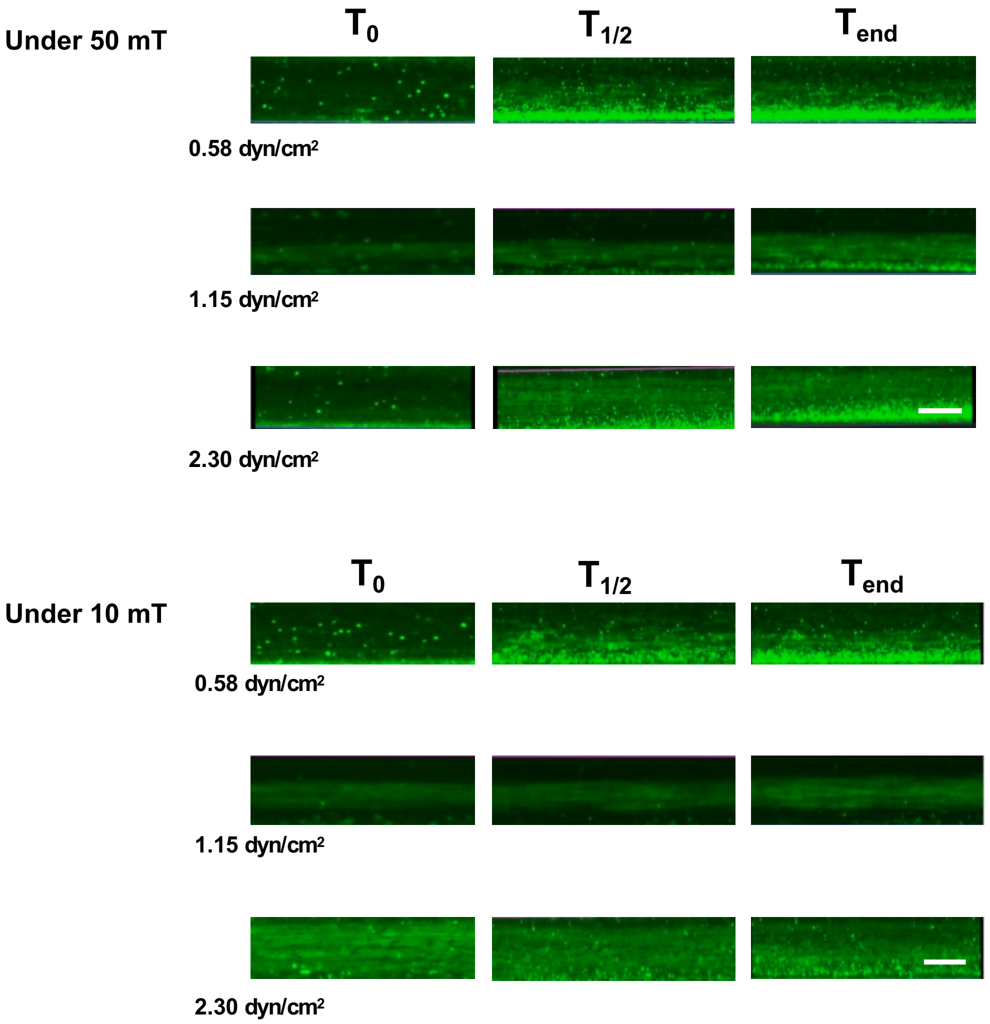


**Fig. S10.** Representative fluorescence time-lapse retention images of immunobots under the application of 50 and 10 mT magnetic field.

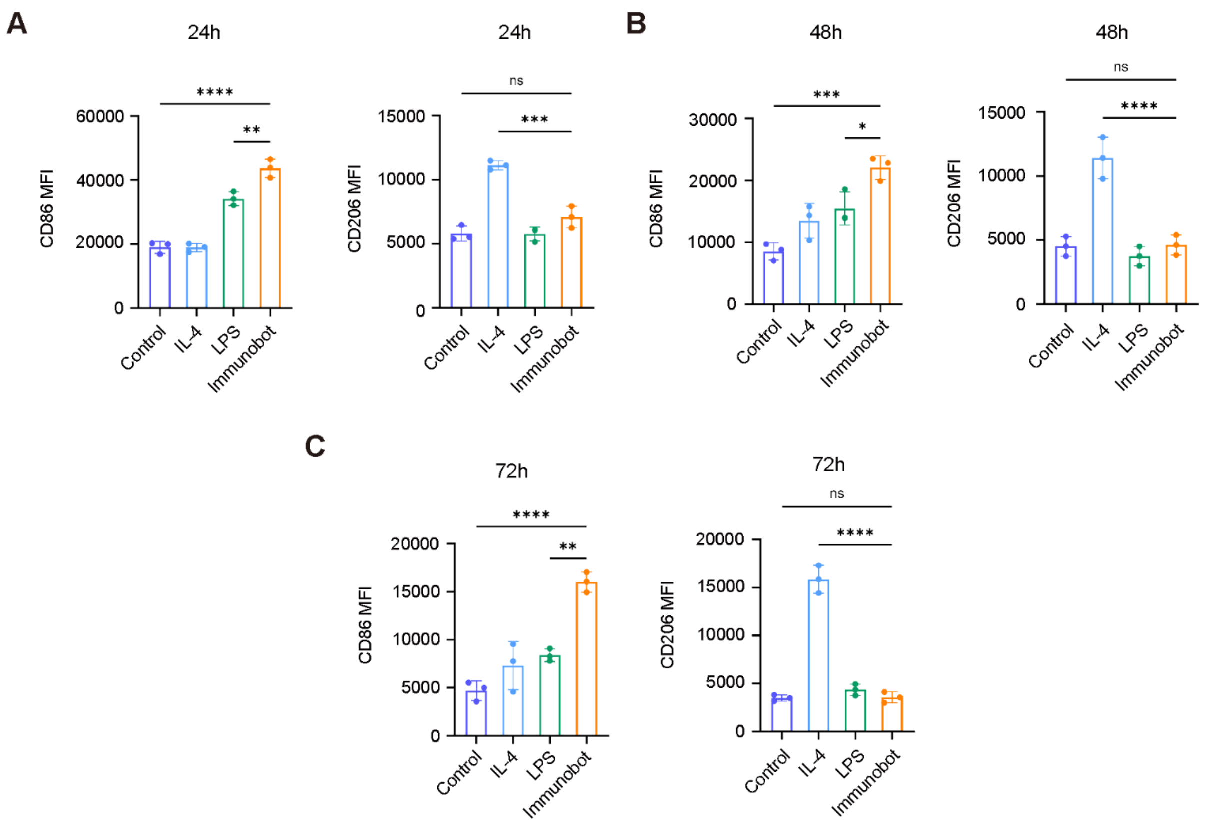


**Fig. S11. Flow cytometric analysis of CD86 and CD206 expression in BMDMs at 24, 48, and 72 h.** BMDMs were treated under the indicated conditions and analyzed by flow cytometry for CD86 and CD206 expression. Quantitative results show the MFI of CD86 and CD206 at 24 h (A), 48 h (B), and 72 h (C). Data are presented as mean ± SD, n = 3. Statistical analysis was performed using one-way ANOVA with Tukey's multiple comparisons test. **P* < 0.05, ***P* < 0.01, ****P* < 0.001, *****P* < 0.0001; ns, not significant.

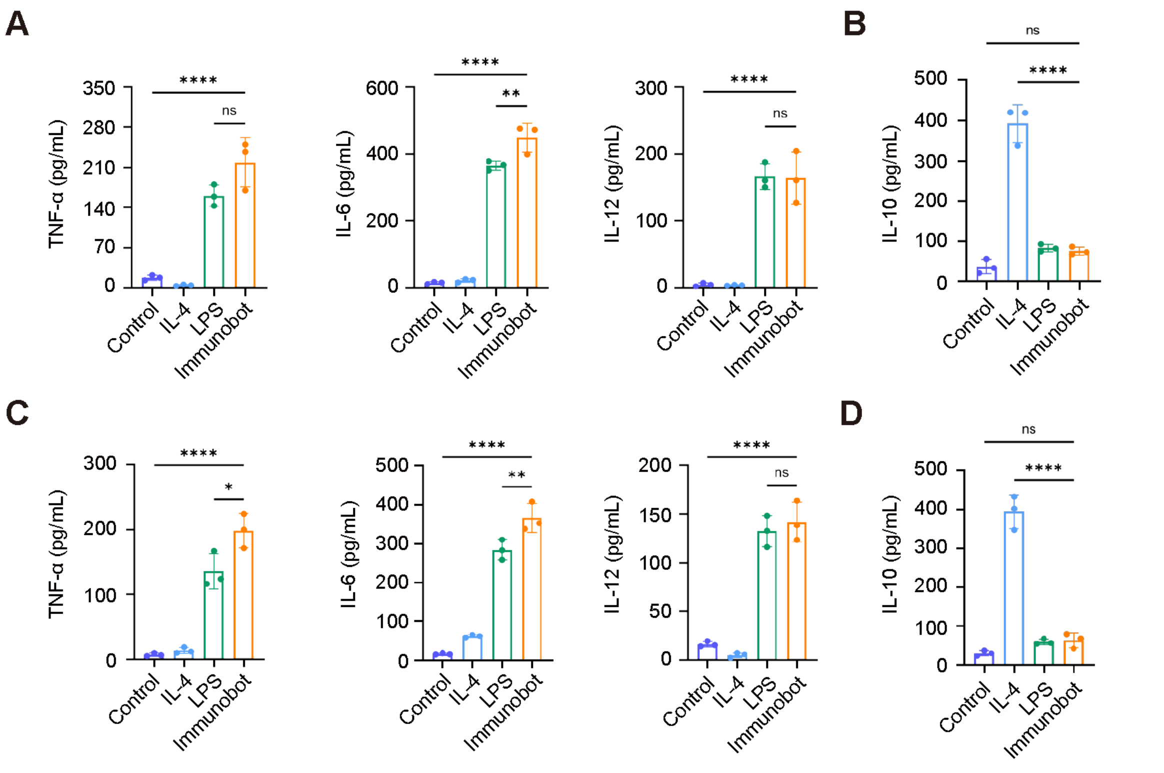


**Fig. S12. ELISA analysis of cytokine secretion in BMDMs under the indicated conditions.** (A) Secretion levels of M1-associated cytokines TNF-α, IL-6, and IL-12 at 48 h. (B) Secretion level of the M2-associated cytokine IL-10 at 48 h. (C) Secretion levels of TNF-α, IL-6, and IL-12 at 72 h. (D) Secretion level of IL-10 at 72 h. Data are presented as mean ± SD (n = 3). Statistical analysis was performed using one-way ANOVA with Tukey's multiple comparisons test. *$P < 0.05$, **$P < 0.01$, ****$P < 0.0001$; ns, not significant.

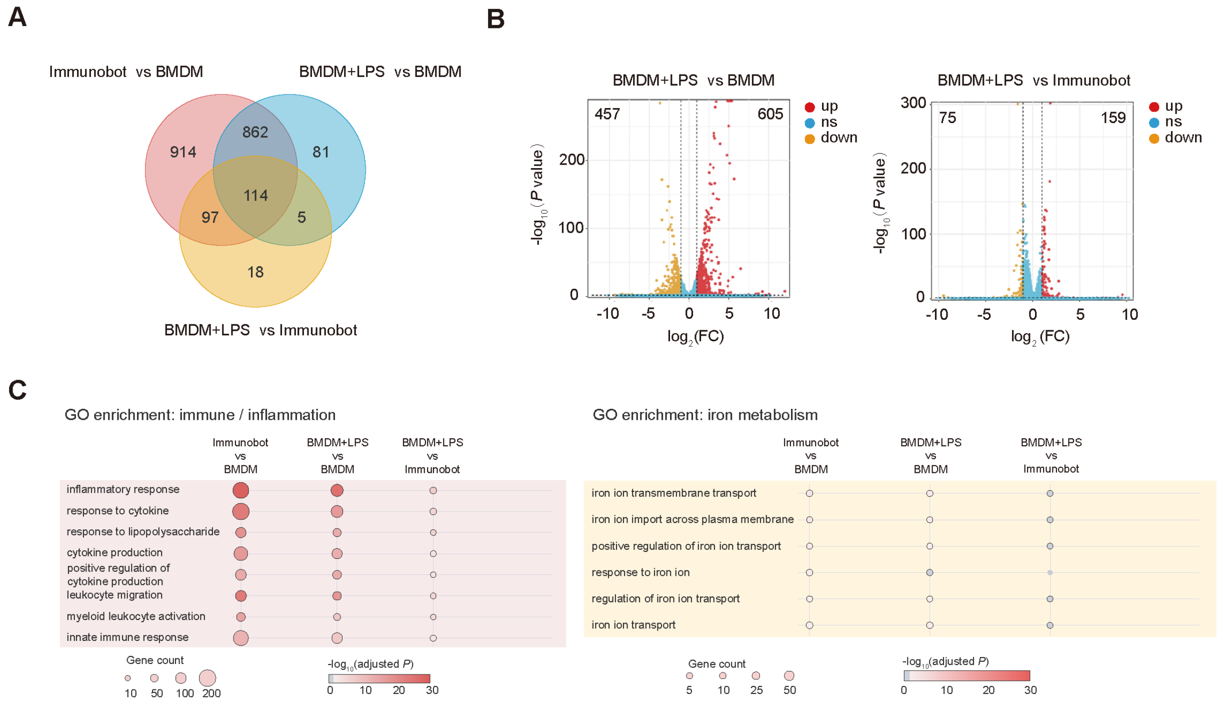


**Fig. S13. RNA-seq profiling of immunobots.** (A) Venn diagram showing shared and unique DEGs among the indicated comparisons. (B) Volcano plots showing DEGs in BMDM + LPS versus BMDM and BMDM + LPS versus immunobot. Red, orange, and blue dots indicate upregulated, downregulated, and non-significant genes, respectively. (C) GO enrichment analysis of immune/inflammatory and iron metabolism-related terms across the indicated comparisons. Bubble size indicates gene count, and color indicates -$\log_{10}$ adjusted *P* value.

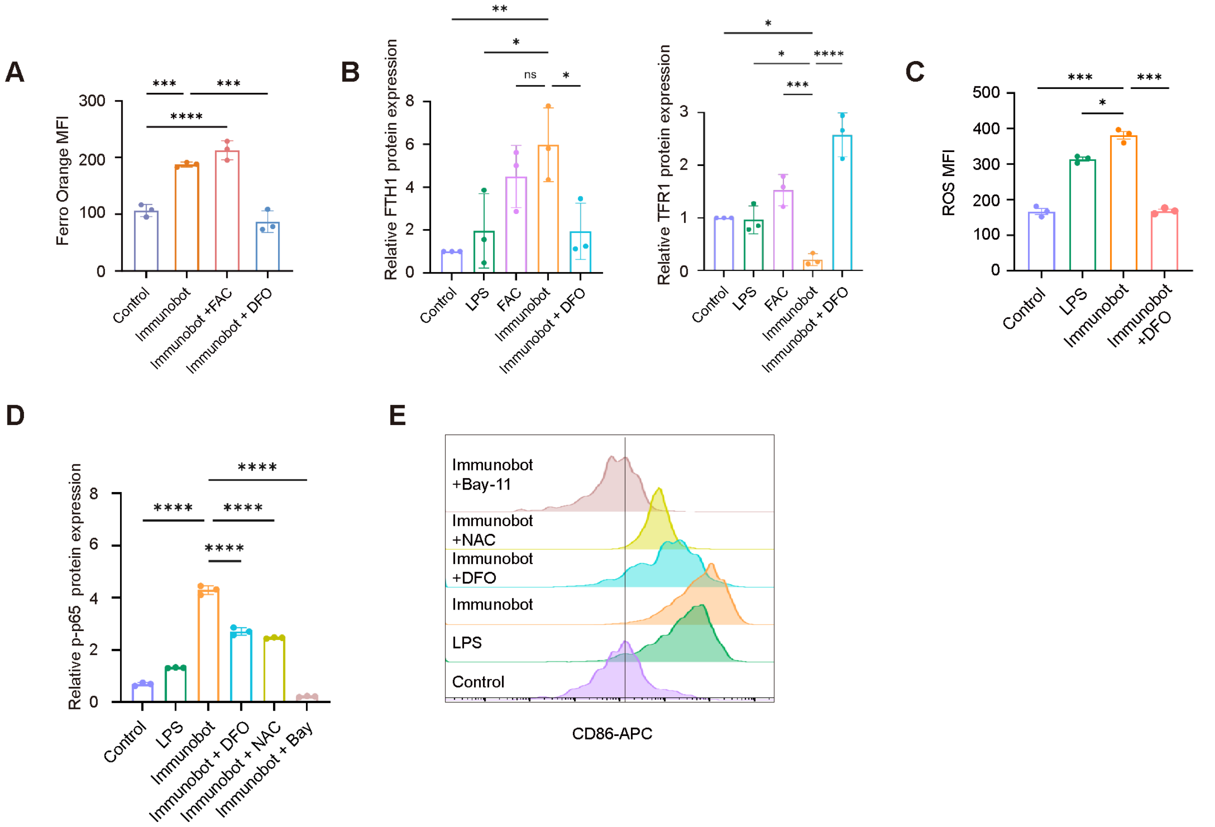


**Fig. S14. Iron-associated activation of immunobots.** (A) Quantification of intracellular labile iron levels by FerroOrange fluorescence intensity in the indicated groups. (B) Densitometric quantification of FTH1 and TFR1 Western blot bands normalized to GAPDH and expressed relative to the control group. (C) Quantification of intracellular ROS levels by fluorescence intensity in the indicated groups. (D) Densitometric quantification of phosphorylated p65 Western blot bands normalized to GAPDH and expressed relative to the control group. (E) Representative flow cytometry histograms showing CD86 expression after treatment. Data are presented as mean ± SD, with dots representing independent biological replicates, n = 3. Statistical significance was determined by one-way ANOVA with Tukey's multiple comparisons test. *$P < 0.05$, **$P < 0.01$, ***$P < 0.001$, and ****$P < 0.0001$; ns, not significant.

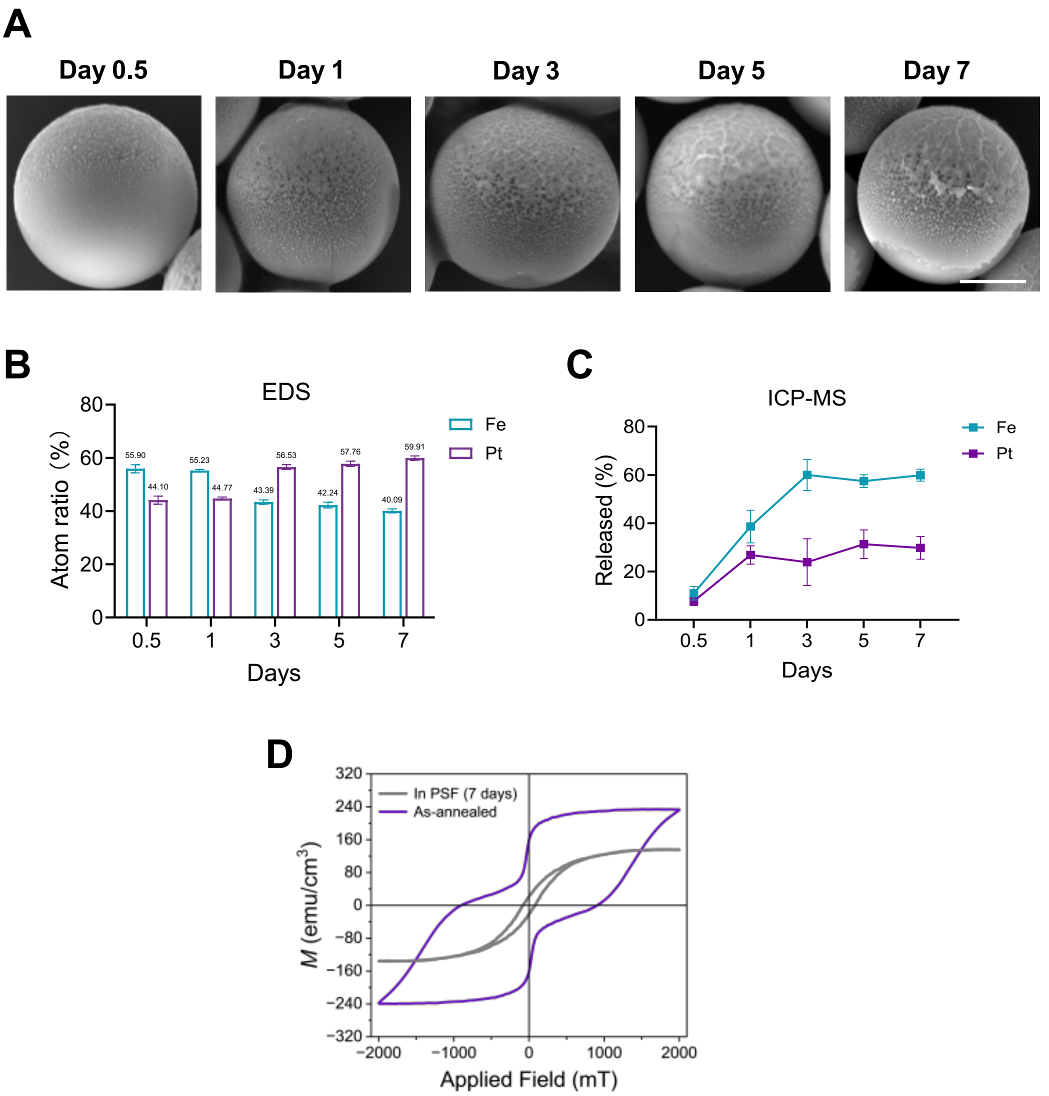


**Fig. S15. Analysis of acid-triggered surface corrosion and preferential Fe leaching from FePt microrollers in phagolysosome simulating fluid (PSF).** FePt microrollers were incubated in acidic PSF (pH ≈ 4.5) at 37 °C for 0.5, 1, 3, 5, and 7 days under shaking. (A) SEM images showing progressive surface roughening and localized pitting/crack-like features on the L10-FePt coating during acid exposure. Scale bar, 2 μm. (B) EDS analysis of the residual coating reveals a time-dependent decrease in Fe atomic percentage and a corresponding increase in Pt atomic percentage. (C) ICP-MS analysis showing the released fractions of Fe and Pt at each time point, defined as the elemental content in the supernatant divided by the total elemental content recovered from the supernatant and pellet. These results indicate partial acid etching of the FePt coating with preferential Fe dissolution and relative Pt enrichment of the remaining surface. (D) VSM hysteresis loop showing the magnetic properties change after 7 days incubation within PSF.

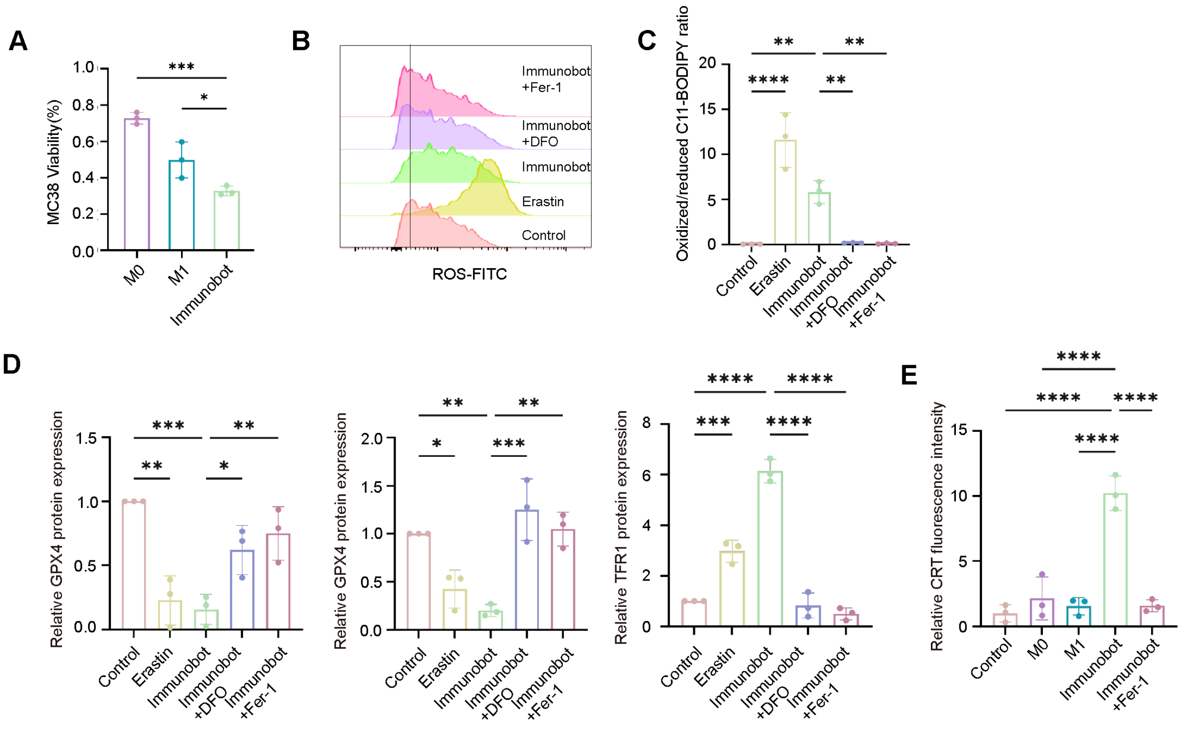


**Fig. S16. Immunobots induce ferroptotic death and immunogenic signaling in MC38 tumor cells.** (A) Viability of MC38 cells after treatment with M0 macrophages, M1 macrophages, or immunobots. (B) Representative flow cytometry histograms showing ROS-FITC fluorescence in MC38 cells treated with control, erastin, immunobots, immunobots plus DFO, or immunobots plus Fer-1. (C) Quantification of lipid peroxidation by the oxidized/reduced C11-BODIPY ratio in MC38 cells under the indicated treatments. (D) Quantification of ferroptosis defense and iron-homeostasis proteins, including GPX4, TFR1, and FTH1, in MC38 cells after the indicated treatments. (E) Quantification of CRT fluorescence intensity in MC38 cells treated with control, M0 macrophages, M1 macrophages, immunobots, or immunobots plus Fer-1. Data are presented as mean ± SD, n = 3 biologically independent samples. Statistical significance was determined by one-way ANOVA with Tukey's multiple-comparisons test. **P* < 0.05, ***P* < 0.01, ****P* < 0.001, and *****P* < 0.0001.

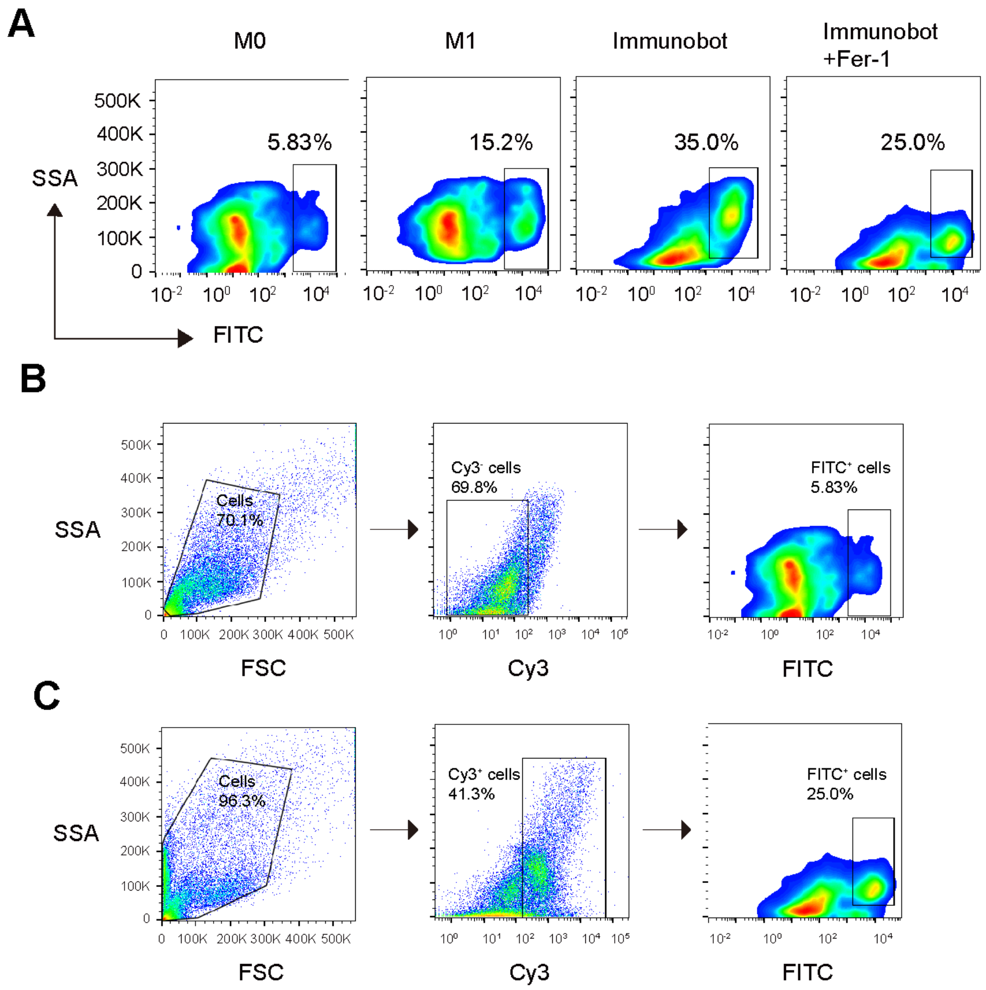


**Fig. S17. Flow cytometric analysis of tumor cell phagocytosis.** (A) Representative flow cytometry plots showing the phagocytosis of tumor cells by M0 macrophages, M1 macrophages, immunobots, and immunobots treated with Fer-1. The percentage of FITC-positive cells represents the phagocytic population. (B, C) Representative gating strategies for M0/M1 macrophages and immunobots, respectively. Cells were gated based on FSC/SSC, followed by Cy3-positive cell selection and FITC-positive cell quantification.

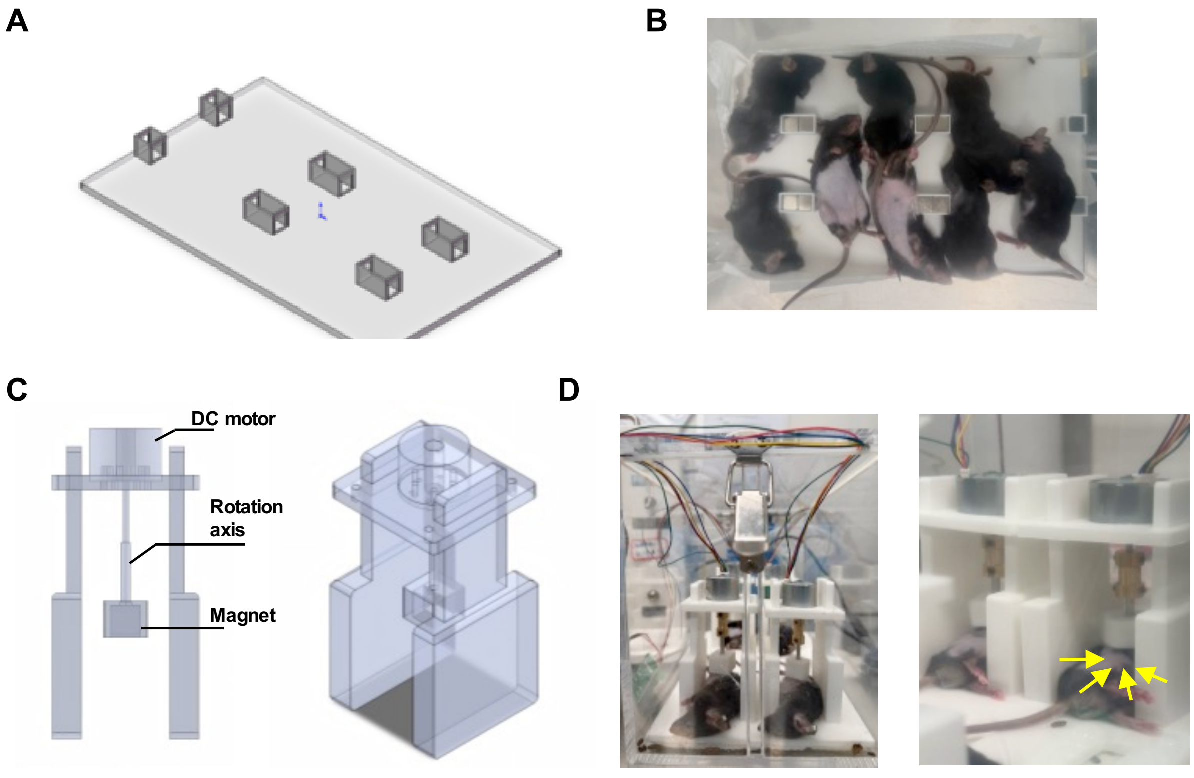


**Fig. S18. *In vivo* magnetic actuation setups.** (A) Permanent magnet fixation model used to apply gradient magnetic field to multiple mice at a time. (B) Representative operation image showing eight mice receiving gradient magnetic field. (C) Computer-aided design of rotating magnetic field application setup. DC motor connected to a permanent magnet. (D) Photograph of RMF application to multiple mice at a time to enhance immunobots penetration.

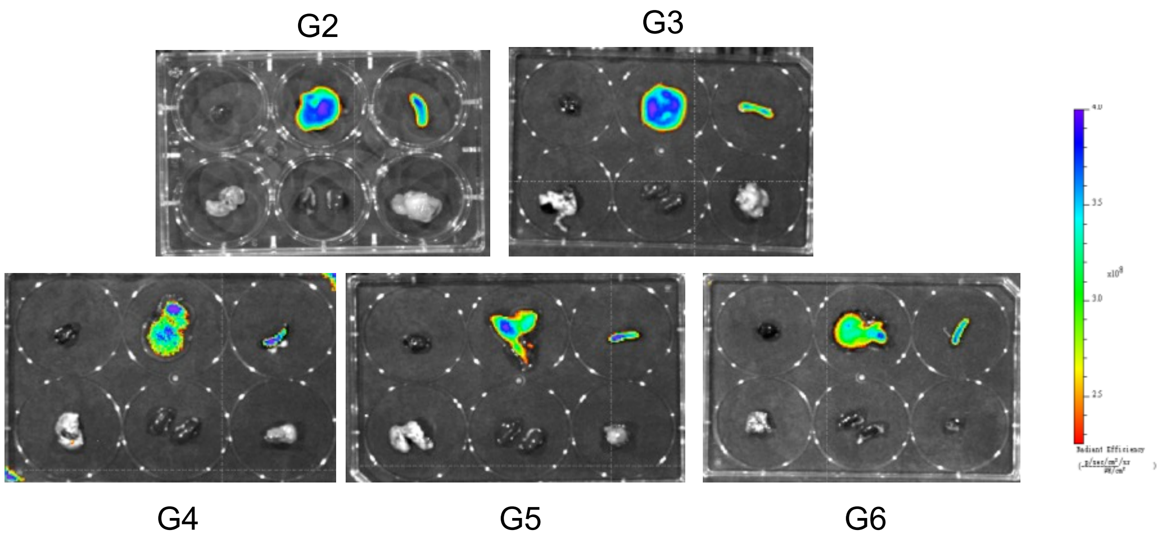


**Fig. S19.** ***Ex vivo*** **biodistribution of DiD-labeled formulations in major organs and tumors at the study endpoint**. In each six-well plate, the top row contains the heart, liver, and spleen, and the bottom row contains the lung, kidney, and tumor.

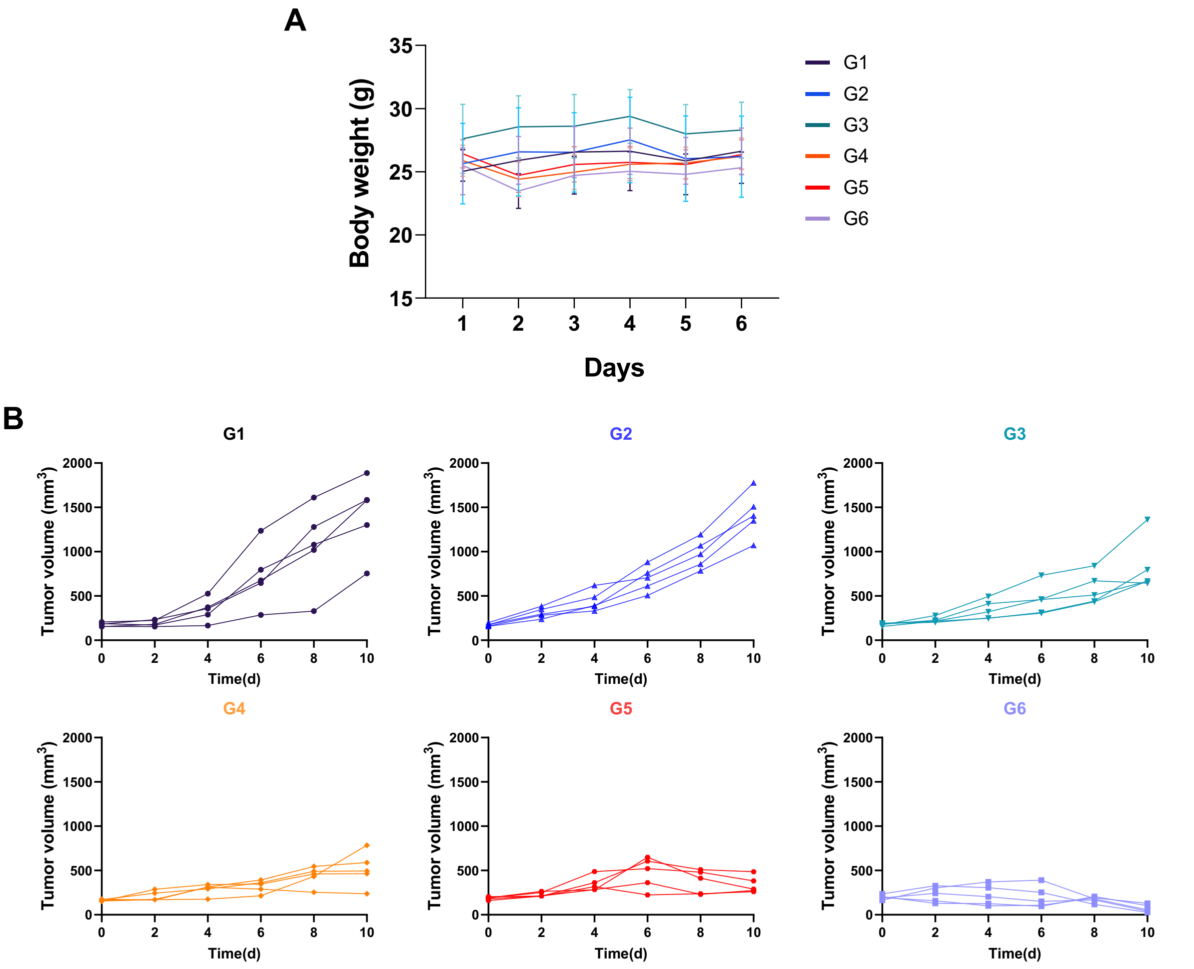


**Fig. S20. Assessment of body weight and normalized tumor growth during treatment.** (A) Changes in mouse body weight across the indicated treatment groups. (B) Tumor growth curves normalized to the initial tumor volume on day 0 across all six groups.

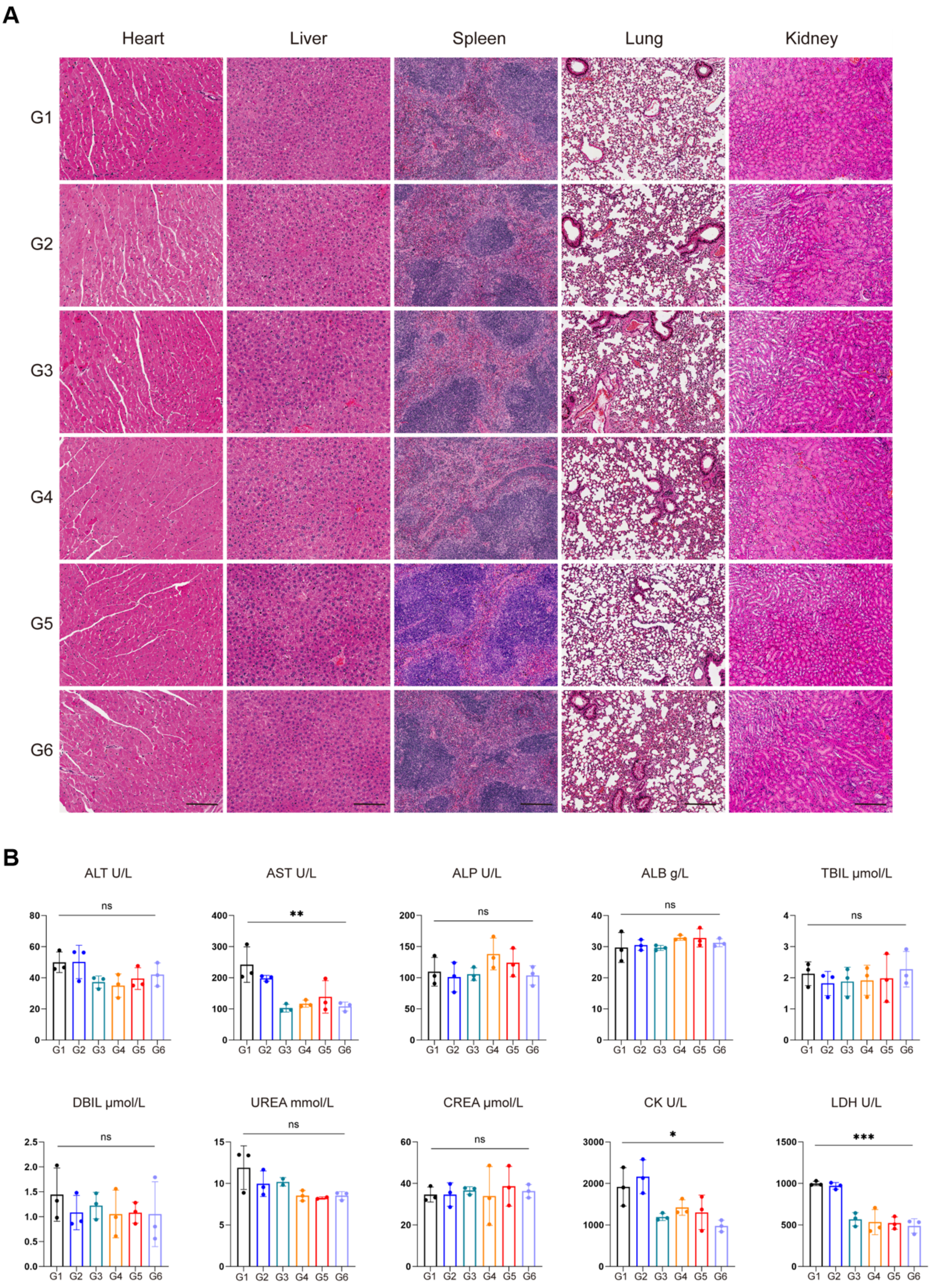


**Fig. S21.** ***In vivo*** **biocompatibility and biosafety evaluation.** (A) Representative H&E-stained sections of major organs (heart, liver, spleen, lung, and kidney) from different treatment groups. Scale bar, 100 μm. (B) Serum biochemical parameters at the sample collection time point in different treatment groups, including ALT, AST, ALP, ALB, TBIL, DBIL, UREA, CREA, CK, and LDH.

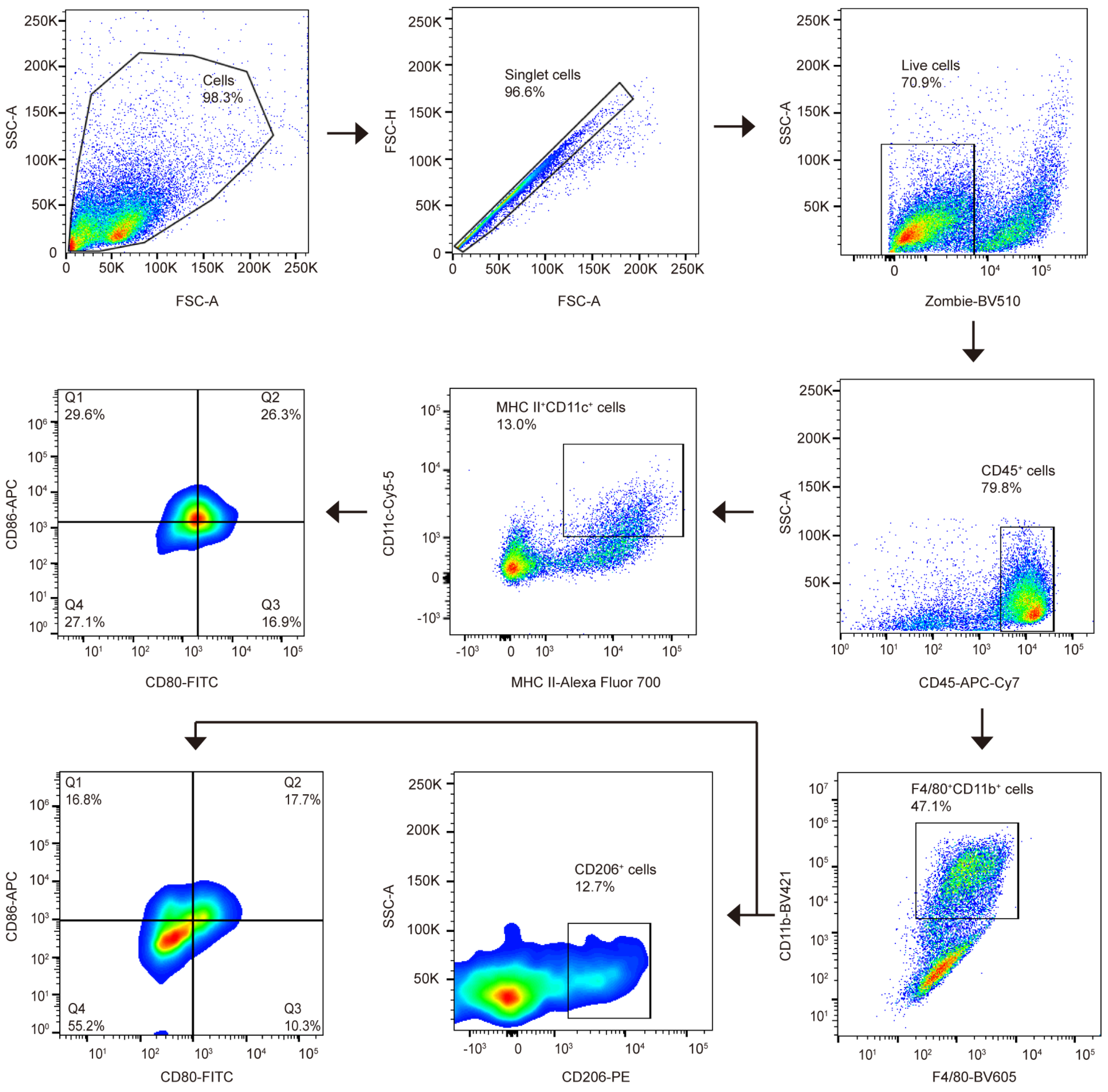


**Fig. S22. Flow cytometry gating strategy for tumor-infiltrating macrophages and dendritic cells (DCs).** Live $CD45^+$ immune cells were gated to identify MHC $II^+CD11c^+$ DCs and $F4/80^+CD11b^+$ macrophages, followed by analysis of CD80/CD86 and CD206 expression.

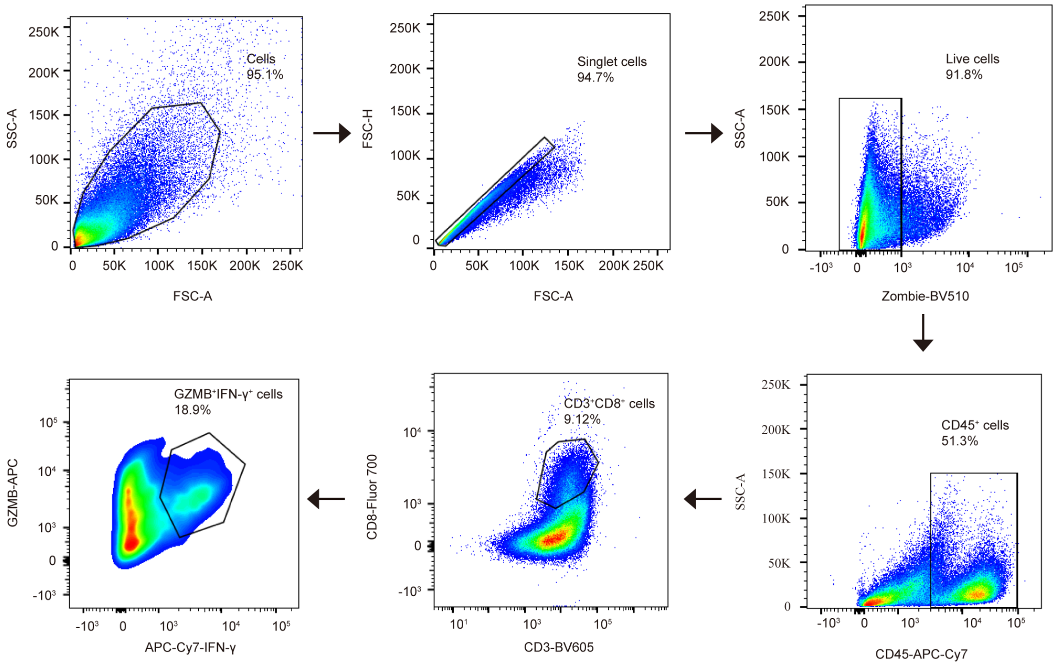


**Fig. S23. Flow cytometry gating strategy for tumor-infiltrating CD8$^{+}$ T cells.** Live CD45$^{+}$ immune cells were gated to identify CD3$^{+}$CD8$^{+}$ T cells, followed by analysis of granzyme B (GZMB) and IFN-γ expression.

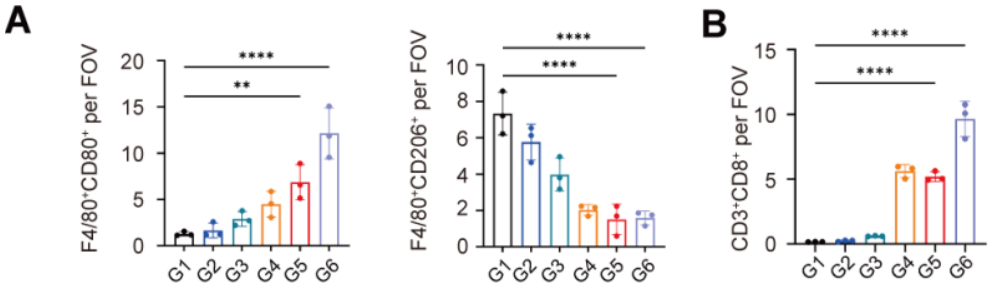


**Fig. S24. Immunofluorescence quantification of macrophage polarization and CD8$^{+}$ T-cell infiltration.** (A) Quantification of F4/80$^{+}$CD80$^{+}$ and F4/80$^{+}$CD206$^{+}$ cells per field of view (FOV) across groups G1 to G6. (B) Quantification of CD3$^{+}$CD8$^{+}$ cells per FOV across groups G1 to G6. Data are presented as mean ± SD; each dot represents one biological replicate. $^{**}P < 0.01$, $^{****}P < 0.0001$.

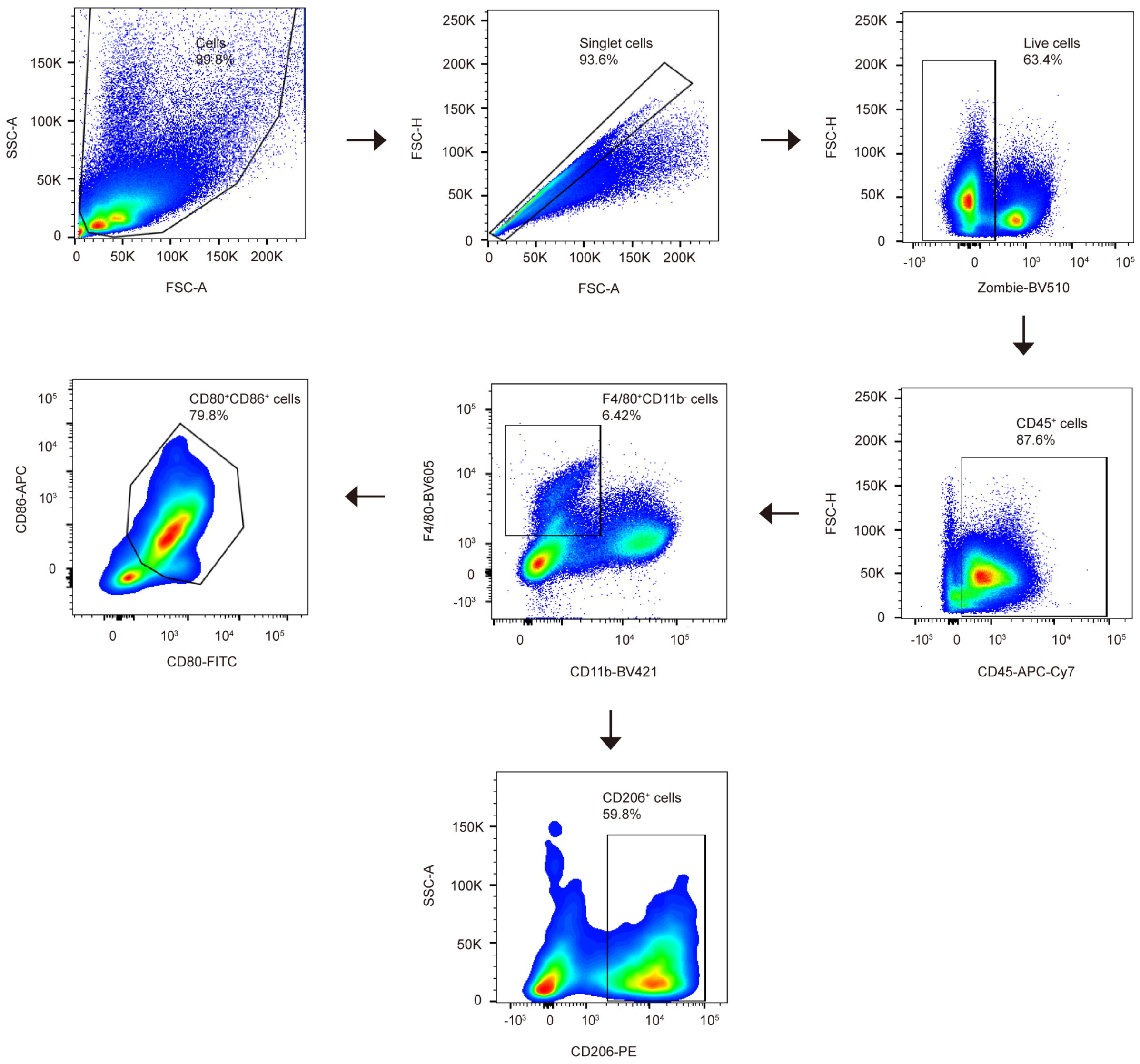


**Fig. S25. Flow cytometric gating strategy for splenic macrophage phenotypes.** Representative gating strategy for the analysis of splenic macrophage phenotypes. Cells were sequentially gated on the basis of FSC-A/SSC-A, singlets, live cells, and CD45$^+$ leukocytes. Splenic macrophages were defined as CD45$^+$F4/80$^+$CD11b$^-$ cells, followed by quantification of CD80$^+$CD86$^+$ M1-like macrophages and CD206$^+$ M2-like macrophages within this population. Percentages indicate representative frequencies within the corresponding parent gates.

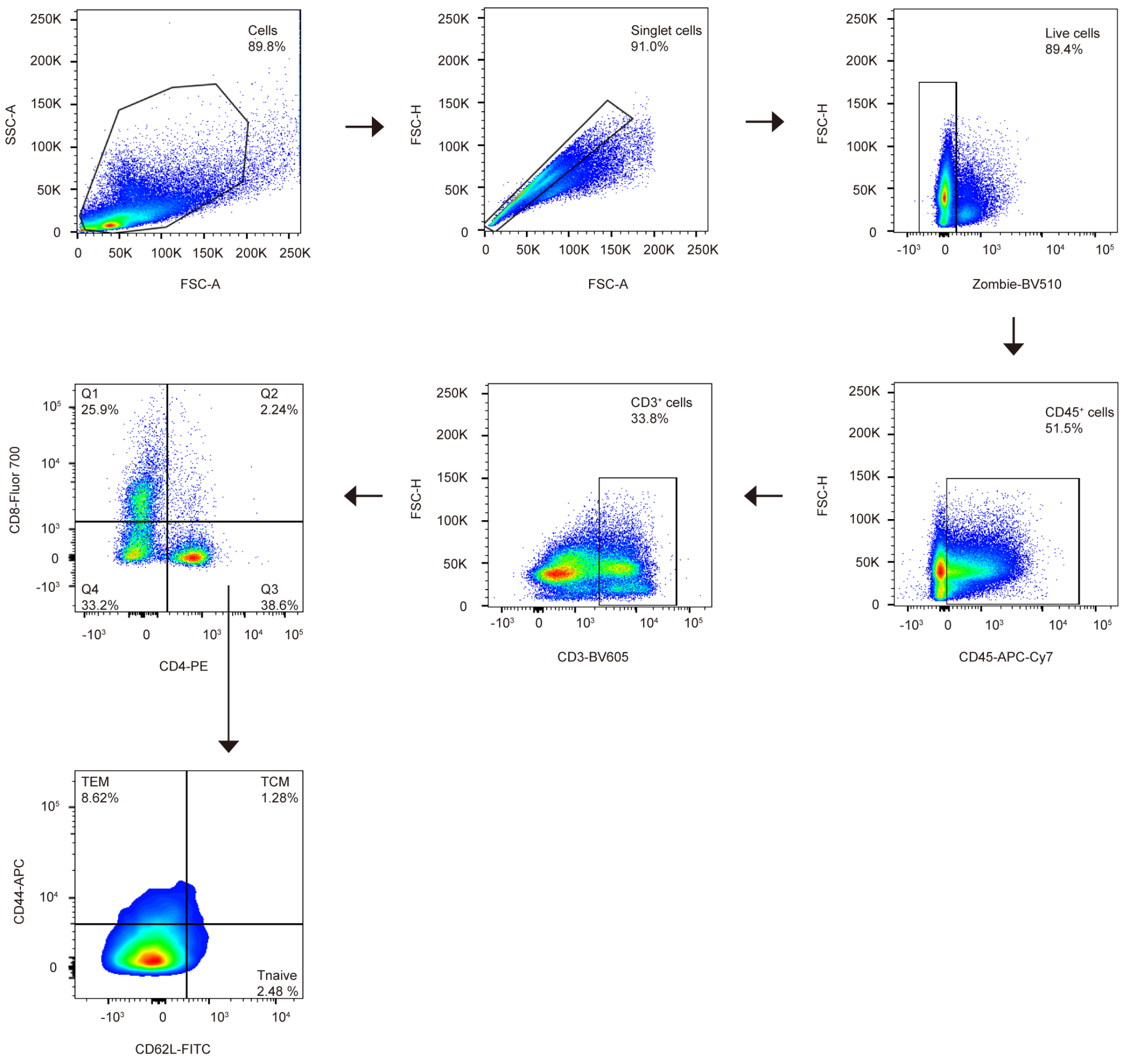


**Fig. S26. Flow cytometry gating strategy for splenic T cells.** Representative gating strategy used to identify splenic T-cell subsets. Cells were sequentially gated by FSC-A/SSC-A, singlets, live cells, and $CD45^+$ leukocytes. T cells were defined as $CD45^+CD3^+$ cells and further analyzed for $CD4^+$ subsets. Memory T-cell phenotypes were determined based on CD44 and CD62L expression, including effector memory T cells (T_EM, $CD44^+CD62L^-$), central memory T cells (T_CM, $CD44^+CD62L^+$), and naïve T cells (T_naive, $CD44^-CD62L^+$). Percentages indicate representative frequencies within the indicated parent gate.

**Table S1. Sputtering power ratio between Fe/Pt targets and corresponding EDS atomic ratio. Data are mean ± SD，n=3.**

| Fe:Pt power ratio | Fe:Pt Atom (%) |
|---|---|
| 200:180 | 45.33 : 55.67 ± 2.18% |
| 200:160 | 51.06 : 49.94 ± 1.35% |
| 200:140 | 55.14 : 45.86 ± 1.42% |
| 200:120 | 58.87 : 42.13 ± 2.27% |

**Table S2. Antibody information used in this study.**

| Antigen | Company | Catalog number | Application |
|---|---|---|---|
| Recombinant Anti-CD8 alpha antibody (Rabbit mAb) | Servicebio | GB15068-50 | Immunofluorescence |
| Anti-F4/80 Rabbit pAb | Servicebio | GB113373-50 | Immunofluorescence |
| Anti-Ki67 Rabbit pAb | Servicebio | GB111141-50 | Immunofluorescence |
| Anti-Mouse Receptor/CD206 Rabbit pAb | Servicebio | GB113497-50 | Immunofluorescence |
| Recombinant Anti-CD86 antibody (Rabbit mAb) | Servicebio | GB150054-50 | Immunofluorescence |
| Anti-CD3 Rabbit pAb | Servicebio | GB11014-50 | Immunofluorescence |
| Anti-CD31 Rabbit pAb | Servicebio | GB113151-50 | Immunofluorescence |
| Recombinant Anti-4-Hydroxynonenal antibody (Mouse mAb) | Servicebio | GB150073-50 | Immunofluorescence |
| Calreticulin Polyclonal antibody | Proteintech | 27298-1-AF | Immunofluorescence |
| APC anti-mouse CD8a | BioLegend | 100711 | Flow cytometry |
| APC/Cyanine7 anti-mouse CD45 | BioLegend | 103115 | Flow cytometry |
| Brilliant Violet 421™ anti-mouse/human CD11b | BioLegend | 101235 | Flow cytometry |
| PE anti-mouse CD4 | BioLegend | 100511 | Flow cytometry |
| Brilliant Violet 605™ anti-mouse F4/80 | BioLegend | 123133 | Flow cytometry |
| PerCP/Cyanine5.5 anti-mouse CD11c | BioLegend | 117327 | Flow cytometry |
| Brilliant Violet 605™ anti-mouse CD3 ε | BioLegend | 100351 | Flow cytometry |
| FITC anti-mouse CD80 | BioLegend | 104705 | Flow cytometry |
| Alexa Fluor® 700 anti-mouse I-A/I-E | BioLegend | 107621 | Flow cytometry |
| APC-conjugated anti-mouse CD86 | BioLegend | 105011 | Flow cytometry |
| PE-conjugated anti-mouse CD206 | BioLegend | 141705 | Flow cytometry |
| APC anti-human/mouse Granzyme B Recombinant | BioLegend | 372204 | Flow cytometry |

| | | | |
|---|---|---|---|
| APC/Cyanine7 anti-mouse IFN-γ | BioLegend | 505849 | Flow cytometry |
| APC anti-mouse/human CD44 | BioLegend | 103011 | Flow cytometry |
| FITC anti-mouse CD62L | BioLegend | 104405 | Flow cytometry |
| Zombie Violet™ Fixable Viability Kit | BioLegend | 423113 | Flow cytometry |
| TruStain FcX™ PLUS (anti-mouse CD16/32) | BioLegend | 156603 | Flow cytometry |
| Ferritin Heavy Chain Polyclonal Antibody | Proteintech | 11682-1-AP | Western blot |
| CD71 Polyclonal Antibody | Proteintech | 10084-2-AP | Western blot |
| GAPDH Monoclonal Antibody | Proteintech | 60004-1-Ig | Western blot |
| Phospho-NF-kappaB p65 (Ser536) (93H1) Rabbit Monoclonal Antibody | Cell Signaling Technology | 3033T | Western blot |
| GPX4 Monoclonal antibody | Proteintech | 67763-1-Ig | Western blot |

**Supplementary Movies**

**Movie S1.**

FePt Janus MR translation speed at different rotating frequency.

**Movie S2.**
Phagocytosis of LMRs by BMDMs.

**Movie S3.**

Comparison of the pre-magnetized and remagnetized immunobots' locomotion performance.

**Movie S4.**

Magnetic navigation of immunobots in diverse environments.

**Movie S5.**

Gradient field capture and flow-resistant retention of immunobots.

**Movie S6.**
*In vivo* magnetic field application system.

**Movie S7.**
Co-localized IVIS and micro-CT imaging of tumor targeting in mice.
